# A JAK/STAT-Pdk1-S6K axis bypasses systemic growth restrictions to promote regeneration

**DOI:** 10.1101/2024.08.05.606658

**Authors:** Ananthakrishnan Vijayakumar Maya, Lena Neuhaus, Liyne Nogay, Aakriti Singh, Lara Heckmann, Isabelle Grass, Jörg Büscher, Katrin Kierdorf, Anne-Kathrin Classen

## Abstract

Inflammation triggers systemic signals that induce growth restrictions in distant organs, a process well characterized in tumor cachexia. While mechanisms allowing tumors to circumvent these systemic growth restrictions have been established, the physiological processes that overcome inflammation-induced growth restrictions during regeneration remain largely unexplored. In our study, we use a model of tissue inflammation and regeneration in developing *Drosophila* imaginal discs to dissect key metabolic and signaling adaptations that, in physiological settings, both induce and overcome systemic growth restrictions. We find that expression of *eiger*, the *Drosophila* TNF-α homolog, induces systemic insulin restriction, as evidenced by reduced expression of *dILP2* and *dILP5*, as well as elevated nuclear dFOXO, and reduced protein translation and proliferation in peripheral tissues. Proliferating cells overcome systemic insulin restriction by upregulating Pdk1, which is both necessary and sufficient to promote phosphorylation of ribosomal protein S6 and protein translation via an Insulin/Akt-independent mechanism. We further demonstrate that JAK/STAT signaling acts upstream to increase Pdk1 levels, delineating a novel JAK/STAT-Pdk1-S6K axis essential for regenerative proliferation. The upregulation of amino acid transporters in the proliferative domain further suggests that regenerating cells preferentially import amino acids, fueling mTORC1 activation. Similar metabolic signatures are observed in a *Drosophila Ras^V12^, scrib* tumor model, suggesting that tumors may co-opt these metabolic pathways to sustain growth in an insulin-restricted environment. Our findings reveal a specialized metabolic program that integrates systemic nutrient mobilization with local metabolic reprogramming, with important implications for understanding of physiological tissue repair but also pathologies such as chronic wounds and cancer.

## Introduction

Tissue damage and inflammation trigger a dynamic interplay between cellular signals and cell behaviors to promote repair and regeneration. Proper orchestration of these responses is crucial, as failure can lead to chronic wound healing pathologies or diseases like cancer (Eming et al, 2017; Huang et al, 2022; MacCarthy-Morrogh & Martin, 2020; Pena & Martin, 2024; Rybinski et al, 2014). To better understand these diseases, a wide range of recent studies aim to dissect the pathological reprogramming of relevant metabolic circuits (Martinez-Reyes & Chandel, 2021; Stine et al, 2022; Wong & Verheyen, 2021). However, a surprising gap exists in our knowledge about the precise metabolic adaptations employed by the normal physiological programs of tissue repair and regeneration (Eming et al, 2021; Eming et al, 2017; Meacham et al, 2022; Zhu & Thompson, 2019). With this study, we aim to provide insight into the local and systemic metabolic adaptations during physiological tissue repair and regeneration.

Central to both metabolism and cellular growth are the Insulin/PI3K/Akt and mTORC1 signaling pathways, which are evolutionarily conserved from invertebrates to vertebrates. Both pathways converge on their shared effector ribosomal protein S6 kinase (S6K), which drives protein synthesis and cellular growth by activating protein translation. The Insulin/PI3K/Akt signaling branch is activated by binding of Insulin to its receptor, stimulating Phosphoinositide-3-kinase (PI3K) to produce Phosphatidylinositol (3,4,5)-trisphosphate (PIP3). PIP3 recruits Phosphoinositide-dependent kinase 1 (Pdk1) and activates Akt, which inhibits nuclear translocation of the transcription factor FOXO. Pdk1 phosphorylates S6K, initiating its activation, whereas optimal S6K activity requires an additional phosphorylation by mTORC1 (Holz et al, 2005; Hopkins et al, 2020; Hoxhaj & Manning, 2020; Manning & Cantley, 2007; Wu et al, 2022). mTORC1 specifically responds to amino acid availability and is therefore central to anabolic growth (Saxton & Sabatini, 2017). While previous studies implicate a role for Insulin/PI3K/Akt and mTORC1 signaling in tissue repair processes, the precise metabolic adaptations remain to be investigated (Kakanj et al, 2016; Nakamura et al, 2020; Ring et al, 2023; Wei et al, 2019).

*Drosophila* models have advanced our understanding of tissue repair, regeneration and metabolism (Fox et al, 2020; Worley & Hariharan, 2022). Specifically, studies in developing imaginal discs or the adult gut have highlighted the role of two key signaling pathways-JNK/AP1 and JAK/STAT-in repair and proliferation. These pathways coordinate a range of responses, from senescent-like cell cycle arrest in damaged cells to compensatory proliferation in adjacent cells (Cosolo et al, 2019; Floc’hlay et al, 2023; Jaiswal et al, 2023; Nakamura et al, 2020; Pena & Martin, 2024; Stevens & Page-McCaw, 2012). Importantly, arrested, JNK-signaling cells produce Unpaired (Upd) cytokines, which activate JAK/STAT signaling in in surrounding cells at the site of inflammatory damage, promoting survival and rapid regenerative proliferation (Crucianelli et al, 2022; Floc’hlay et al, 2023; Herrera & Bach, 2019; Herrera et al, 2013; Jaiswal et al, 2023; La Fortezza et al, 2016; Santabarbara-Ruiz et al, 2015; Sustar et al, 2011; Worley et al, 2022). The distinct functional demands of senescent and rapidly proliferating cells raise the question about how these distinct cell populations metabolically adapt to successfully support tissue repair.

Tissue repair and tumor development share striking similarities; in fact, tumors have long been compared to non-healing wounds (Dvorak, 1986). Accordingly, *Drosophila* tumor models activate JNK/AP1 and JAK/STAT signaling, which promote tumor progression (Enomoto & Igaki, 2022; Herrera & Bach, 2019; La Marca & Richardson, 2020). To support their growth, tumors secrete signaling molecules that initiate inter-organ signaling and systemic metabolic responses (Bilder et al, 2021; Hodgson et al, 2021; Liu et al, 2022). For instance, the Insulin-like peptide 8 (Dilp8), when secreted by tumors, disrupts hormone balance by reducing Ecdysone and Insulin production through direct effects on the ring gland and Insulin-producing cells (IPCs), with the effect of halting developmental progression of the tumor-bearing host (Colombani et al, 2012; Garelli et al, 2012). Other cytokines, such as Ecdysone-inducible gene L2 (ImpL2), or the TNFα homologue Eiger (Egr), directly or indirectly trigger lipolysis and proteolysis to promote nutrient release from muscles as well as the fat body, an adipose tissue central for nutrient storage and energy homeostasis (Agrawal et al, 2016; Ding et al, 2021; Figueroa-Clarevega & Bilder, 2015; Gutierrez et al, 2007; Hodgson et al, 2021; Lodge et al, 2021; Rajan & Perrimon, 2012; Romao et al, 2021; Song et al, 2019). Amino acids or sugars are released in this manner and subsequently absorbed by tumors and facilitate their anabolic growth (Cong et al, 2021; Khezri et al, 2021; Newton et al, 2020). This inter-organ signaling network and metabolic state resembles cachexia, a clinical syndrome characterized by weight loss, muscle atrophy and fatigue, typically observed in chronic inflammatory conditions, including cancer (Argiles et al, 2014; Bilder et al, 2021; Hodgson et al, 2021; Liu et al, 2022; Setiawan et al, 2023). While these studies reveal oncogenic metabolic signaling networks, the metabolic signaling networks employed during physiological tissue repair and regeneration remain poorly understood. Previous studies demonstrate that systemically acting cytokines may also be released upon tissue damage in the absence of oncogenic transformation (Colombani et al, 2012; Garelli et al, 2012; Gontijo & Garelli, 2018; Hackney & Cherbas, 2014; Halme et al, 2010; Sun & Poss, 2023), and fat body break-down changing Methionine, S-adenosylmethionine and Kynurenine metabolism promotes imaginal disc regeneration (Kashio & Miura, 2020; Kashio et al, 2016). However, the integration of local and systemic metabolic adaptations promoting physiological repair and regeneration remain poorly characterized. In our study we combine genetic analysis, quantitative imaging, untargeted metabolomics and information from single-cell RNA sequencing data to outline the local and systemic adaptations that selectively support fast-proliferating cells during regeneration through an Insulin-independent JAK/STAT-Pdk1-S6K signaling axis.

## Results

### (1) Proliferating cells maintain high levels of protein translation during regeneration

To induce regeneration, we expressed the *Drosophila* homologue of TNF-α, known as Eiger (Egr), for a 24 h period within the imaginal wing pouch using the *rn-GAL4* driver (Smith-Bolton et al, 2009). As expected, *egr-*expression caused significant cell death, accompanied by the activation of the JNK/AP-1 activity reporter TRE-RFP (**Fig 1A-C; Fig S1A,B**) (Chatterjee & Bohmann, 2012; Cosolo et al, 2019; Jaiswal et al, 2023; La Fortezza et al, 2016). The central disc region with high JNK signaling exhibited markers of cellular senescence, including increased senescence-associated β-galactosidase activity (**Fig. S1C,D**), upregulation of the matrix metalloprotease MMP-1 and cytokines of the Upd family, as well as a JNK-signaling induced cell cycle arrest in G2 (Cosolo et al, 2019; Floc’hlay et al, 2023; Harris et al, 2020; Jaiswal et al, 2023; Worley et al, 2018; Worley et al, 2022). In contrast, cells within 40 µm surrounding this central JNK signaling domain were highly proliferative, as detected by EdU incorporation (**Fig 1A,B,D**) (Crucianelli et al, 2022; Smith-Bolton et al, 2009). To facilitate quantification of cell behaviors and account for disc size variation, we defined a conservative 20µm band outside the high JNK signaling domain as the ‘proliferative domain’ (PD^egr^) for the remainder of this study.

**Figure 1.**
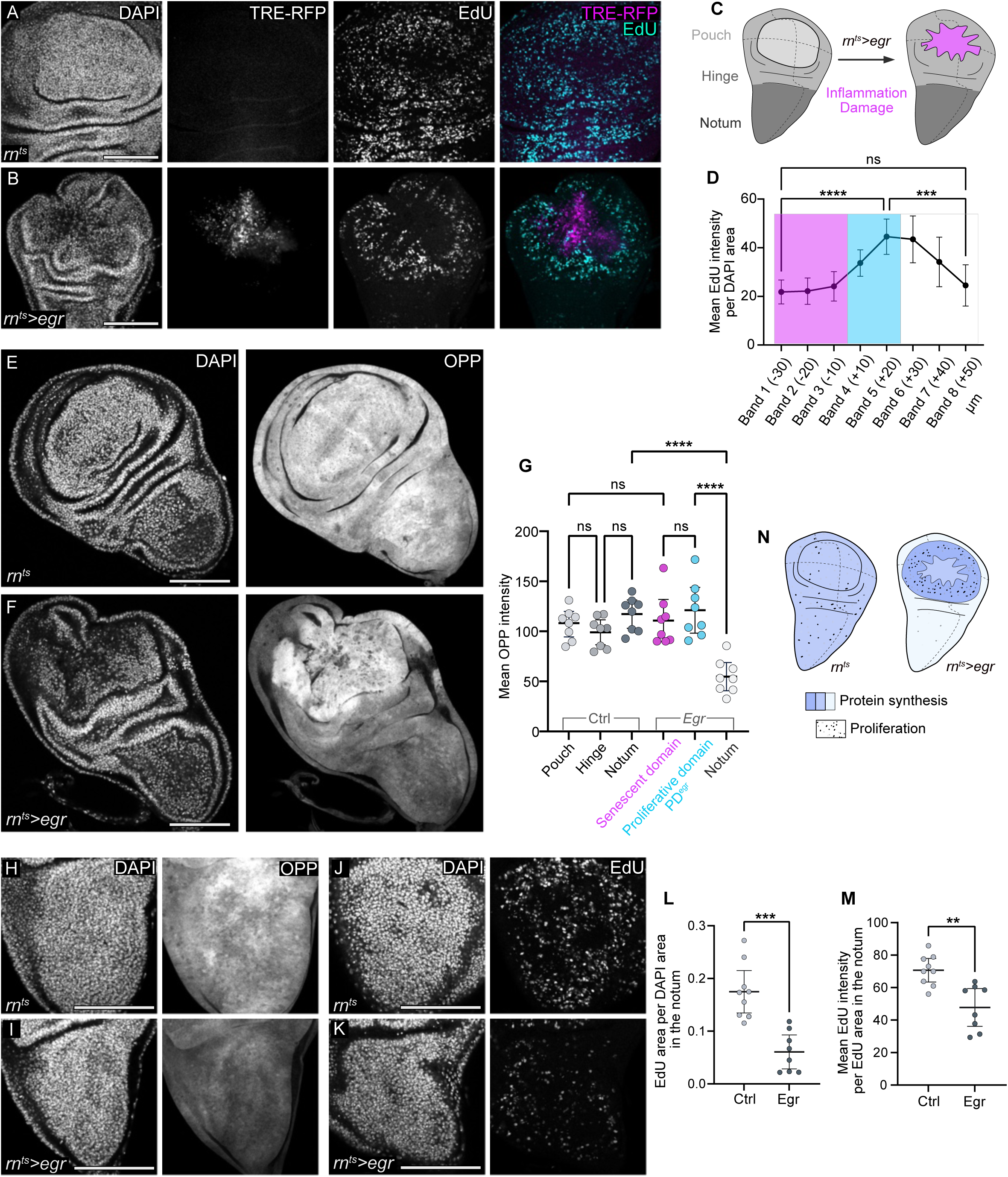
Spatial organization of cell proliferation and protein synthesis induced by inflammatory damage in wing imaginal disc. **A,B.** Control wing disc (A) and a genetically ablated wing disc (B) after 24 hours of *egr-*expression in the wing pouch domain (see Fig. 1C) under the control of the *rn*-GAL4 driver (*rn^ts^* and *rn^ts^>egr*, respectively). TRE-RFP visualizes JNK-pathway activity (magenta or grey), and EdU incorporation visualizes DNA replication (cyan or grey). Discs were stained with DAPI to visualize nuclei. **C.** Schematic representation of a third instar wing imaginal disc, highlighting different regions (pouch, notum, and hinge) on the left. On the right, schematic representation of a wing imaginal disc after 24 hours of *egr-*expression in the pouch. The magenta area identifies the JNK-signaling domain representing inflammatory tissue damage. **D.** Quantification of average EdU intensity in the DAPI area contained within 10µm bands segmented inward and outward from the edge of the high JNK-signaling domain in *egr-*expressing discs. The high JNK-signaling domain is shaded in magenta, the proliferative domain marking the regenerating region is shaded in cyan. Mean and 95% CI (confidence interval) are shown. Statistical significance was tested using a one-way ANOVA followed by Tukey’s post-hoc test for multiple comparisons (n=7). P-values: Band 1 vs Band 5 = 0.0001; Band 5 vs Band 8 = 0.0004; Band 1 vs Band 8 = 0.9984 (ns) **E,F.** Protein synthesis visualized by O-proparyl-puromycin (OPP) incorporation into newly synthesized proteins in a control wing disc (E) and an *egr-*expressing disc (F). Discs were stained with DAPI to visualize nuclei. **G.** Quantification of mean OPP intensity in three different regions of control and *egr-*expressing discs. In control disc, OPP intensity was measured in the pouch, hinge, and notum, while in *egr-*expressing discs, OPP intensity was measured in the high JNK signaling domain (approximating the central pouch), the proliferating region (approximating the peripheral pouch and hinge), and the notum. Mean and 95% CI are shown. Statistical significance was tested using a one-way ANOVA followed by Tukey’s post-hoc test for multiple comparisons (control: n=8, *egr-*expressing disc: n=8). P-values: Proliferative region vs Quiescent notum < 0.0001; Notum vs Quiescent notum < 0.0001; Pouch vs JNK domain = 0.9992 (ns). **H, I.** Protein synthesis visualized by OPP incorporation in the notum of control (H) and *egr-*expressing discs (I). **J,K.** EdU incorporation to visualize DNA replication in the notum of control (J) and *egr-*expressing discs (K). **L.** EdU area per DAPI area quantified in the notum of control and *egr-*expressing discs, serving as a proxy for the number of cells undergoing DNA replication. Mean and 95% CI are shown. Statistical significance was tested using a two-tailed Unpaired t-test (control: n=9, *egr-*expressing disc: n=8), with a p-value = 0.0001. **M.** Mean EdU intensity per EdU-positive area quantified in the notum of control and *egr-*expressing discs, serving as a proxy for DNA replication speed. Mean and 95% CI are shown. Statistical significance was tested using a two-tailed Unpaired t-test (control: n=9, *egr-*expressing disc: n=8), with a p-value = 0.0011. **N.** Schematic representation of protein synthesis rates (blue shades) and mitotic activity (black dots) in control disc (left) and *egr-*expressing discs (right). Scale bars: 100 μm. Fluorescence intensities are reported as arbitrary units.

How do these proliferating cells meet their metabolic needs? To answer this question, we monitored protein translation using O-propargyl-puromycin (OPP)-incorporation assays (Aviner, 2020). This assay revealed uniform levels of protein synthesis throughout control wing imaginal discs. In *egr-*expressing discs, protein synthesis in the proliferative and the JNK signaling domains proceeded at levels similar to control discs (**Fig 1E-G**). Interestingly, the notum of *egr-*expressing discs exhibited a significant decrease in OPP incorporation, which correlated with a marked reduction in EdU incorporation (**Fig 1G-M**). The contrasting levels of protein synthesis, proliferation and signaling between different regions reveal that inflammatory damage induces at least three distinct cell populations with different cellular programs: 1) a senescent cell population exhibiting high protein synthesis and JNK/AP-1 activity; 2) an cycling cell population exhibiting high protein synthesis and low JNK/AP-1 activation; and 3) notum cells exhibiting low protein synthesis, low JNK/AP-1 activity and slow cell cycling (**Fig 1N**). While these observations mirror proliferation dynamics visualized in earlier studies (Smith-Bolton et al, 2009; Sustar et al, 2011; Sustar & Schubiger, 2005), we wanted to better understand how these differences reflect a need to integrate local and systemic metabolic demands during regeneration.

### (2) Tissue damage induces systemic insulin restriction

A reduction in protein translation rates in peripheral tissue domains like the notum suggested that insulin signaling, normally supporting protein translation through S6K activation, is reduced. Notably, cells with active JNK signaling express high levels of Dilp8, a known antagonists of insulin production by insulin-producing cells (IPCs) in the larval brain (**Fig 2A,B**) (Garelli et al, 2012; Vallejo et al, 2015). Expression of *eiger* could thus limit anabolic growth despite nutrient intake by feeding (Agrawal et al, 2016; Figueroa-Clarevega & Bilder, 2015; Gontijo & Garelli, 2018; Ingaramo et al, 2020; Rajan & Perrimon, 2012; Vallejo et al, 2015; Wang et al, 2005). To determine if *egr-*expressing larvae indeed restrict Insulin expression, we analyzed the expression of the *Drosophila* Insulin-like peptides *dILP2* and *dILP5* (Semaniuk et al, 2020). We found that *dILP2* and *dILP5 expression* was significantly reduced in *egr-*expressing larvae, and approached levels seen in larvae starved for 24 h (**Fig 2C**), a condition known to reduce insulin expression due nutrient limitation (Semaniuk et al, 2020; Sudhakar et al, 2020).

**Figure 2.**
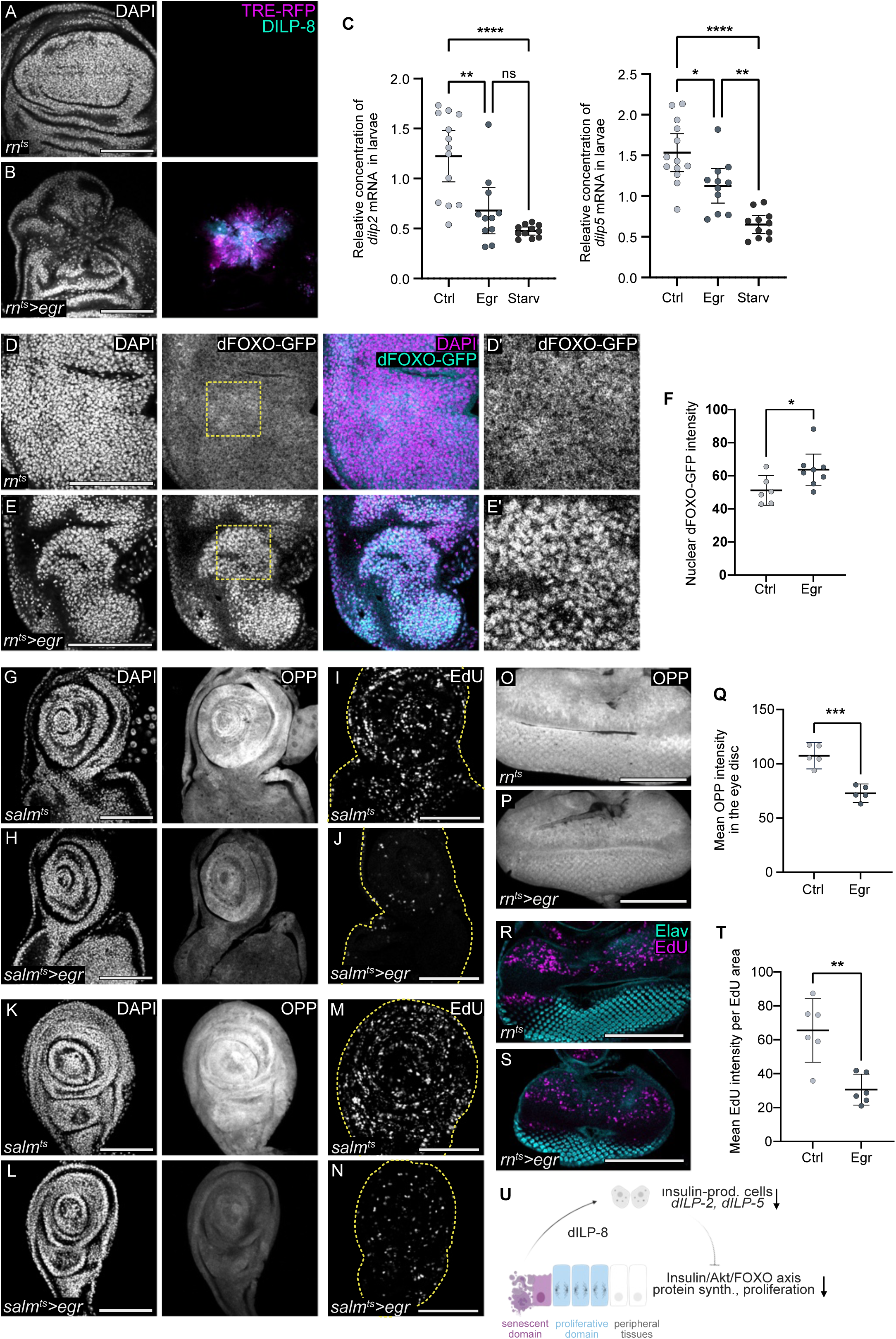
Inflammatory tissue damage induces Insulin restriction and a systemic reduction in growth and cell proliferation. **A, B.** Expression of Dilp8-GFP (cyan) in control (A) and *egr-*expressing discs (B). TRE-RFP visualizes JNK-pathway activity (magenta). Discs were stained with DAPI to visualize nuclei. dFOXO-GFP expression is visualized using BDSC 38644 fly line. **C**. Levels of *dilp2* and *dilp5* mRNA relative to *Rpl1* mRNA were quantified in control, *egr-*expressing, and starved male larvae using quantitative PCR (qPCR). Day6 larvae were used for all conditions. Both genotypes, except starved larvae, undergo temperature shift for 24 hours. For experimental larvae *egr* expression was induced for 24h in the wing pouch using *rn*-GAL4 driver, while starved larvae underwent 24 h starvation. Mean and 95% CI are shown. Statistical significance was tested using the Kruskal-Wallis test followed by Dunn’s multiple comparison test (control larvae: n=13, *egr-*expressing larvae: n=11, starved larvae: n=11). P-values for *dilp2*: Control vs egr = 0.0075; Control vs Starved < 0.0001; egr vs Starved = 0.1138 (ns). P-values for *dilp5*: Control vs egr = 0.0440; Control vs Starved < 0.0001; egr vs Starved = 0.0073. **D, E.** Expression of dFOXO-GFP (cyan and grey) in the notum of control (D) and *egr-*expressing wing imaginal discs (E). Both genotypes were temperature-shifted for 24hours; in panel E, egr expression is restricted to the rotund domain of the wing disc. Discs were stained with DAPI (magenta and grey) to visualize nuclei. High nuclear translocation of dFOXO indicates low insulin signaling. BDSC: 38644 fly line was used. D’ and E’ show magnified view of the regions indicated by the yellow dotted boxes in panels D and E respectively. **F**. Quantification of mean nuclear dFOXO-GFP intensity in the notum of control (D) and *egr-*expressing wing imaginal discs (E). Mean and 95% CI are shown. Statistical significance was tested using a two-tailed Unpaired t-test, p-value = 0.0427 (control: n=6, *egr-*expressing disc: n=8). **G, H.** Protein synthesis visualized by OPP incorporation in the eye-antennal imaginal disc, dissected from larvae with control (G) or *egr-*expressing (H) wing imaginal discs using *salm-*GAL4 driver. The expression of *salm-GAL4* is localized to the central pouch. **I, J.** EdU incorporation to visualizes DNA replication in eye-antennal imaginal discs, dissected from larvae with control (I) or *egr-*expressing (J) wing imaginal discs using *salm-*GAL4 driver. The dotted yellow line outlines the eye-antennal disc. **K, L.** Protein synthesis visualized by OPP incorporation in the leg imaginal disc, dissected from larvae with control (K) or *egr-*expressing (L) wing imaginal discs using *salm-*GAL4 driver. **M, N.** EdU incorporation to visualizes DNA replication in leg imaginal discs, dissected from larvae with control (M) or *egr-*expressing (N) wing imaginal discs using *salm*-GAL4 driver. The dotted yellow line outlines the leg disc. **O, P.** Protein synthesis visualized by OPP incorporation in the eye imaginal disc, dissected from larvae with control (O) or *egr-*expressing (P) wing imaginal discs using *rn*-GAL4 driver. **Q.** Quantification of mean OPP intensity in eye imaginal discs, dissected from larvae with control or *egr-*expressing wing imaginal discs. Mean and 95% CI are shown. Statistical significance was tested using a two-tailed Unpaired t-test, p-value = 0.0002 (control: n=5, *egr-*expressing disc: n=5). **R, S.** EdU incorporation (magenta) to visualizes DNA replication in eye imaginal discs, dissected from larvae with control (R) or *egr-*expressing (S) wing imaginal discs using *rn*-GAL4 driver. Eye disc differentiating photoreceptors are marked with Elav staining (cyan). **T.** Quantification of mean EdU intensity per EdU-positive area serving as a proxy for DNA replication speed, quantified in the eye imaginal discs dissected from larvae with control or *egr-*expressing wing imaginal discs. Mean and 95% CI are shown. Statistical significance was tested using the two-tailed Unpaired t-test, p-value = 0.0015 (control: n=6, *egr-*expressing disc: n=6). **U**. A model illustrating expression of dILP-8 from senescent-like cells in the high JNK signaling domain. DILP-8 causes the downregulation of insulin-like peptides (dILP-2 and dILP-5) from insulin producing cells (IPCs) in the larval brain. This results in a systemic reduction of Insulin/Akt/FOXO signaling, leading to reduced protein synthesis and proliferation in peripheral tissues. Scale bars: 100 μm. Fluorescence intensities are reported as arbitrary units.

To understand if this reduction in *dILP2* and *dILP5* expression correlates with reduced Insulin signaling in peripheral tissues, we examined nuclear localization of the dFOXO, a key downstream effector of canonical Insulin signaling (Hoxhaj & Manning, 2020; Manning & Toker, 2017). We observed elevated nuclear dFOXO in the notum of *egr-*expressing discs (**Fig 2D-F**), demonstrating that reduced *dILP2* and *dILP5* expression correlated with systemic attenuation of insulin signaling. This attenuation would explain the low rates of protein translation observed in the notum. In support of this conclusion, we found that low rates of protein translation in nota did not correlate with activation of JNK stress signaling or apoptosis, indicating that low Insulin signaling likely causes the observed reduction in protein synthesis (**Fig S2A,B**). Importantly, this systemic reduction in protein translation was also evident in other imaginal discs, such as the leg and the eye, and this effect was robustly reproduced by *eiger* expressed under the more restricted spatial pattern of *salm-GAL4* (**Fig 2G-T, Fig S2C-K**). In all cases, the reduction in translational capacity correlated with decreased proliferation, with the developing eye showing reduced EdU incorporation overall and in the second mitotic wave, specifically (**Fig S2H,I**). Taken together, these findings reveal a widespread decline in cell proliferation and protein synthesis in *egr-*expressing larvae, which correlates with restricted Insulin production and signaling (**Fig 2U**). These observations are consistent with previous reports of reduced disc sizes following other types of tissues damage and resemble systemic changes induced by inflammatory tumors (Boulan et al, 2019; Ding et al, 2021; Figueroa-Clarevega & Bilder, 2015; Garelli et al, 2012; Hodgson et al, 2021; Jiang et al, 2022; Khezri et al, 2021; Lodge et al, 2021; Smith-Bolton et al, 2009; Song et al, 2019; Sustar & Schubiger, 2005).

### (3) Regenerative proliferation is supported by fat body catabolism, amino acid importers and TOR signaling

How can the proliferative domain maintain high rates of protein translation and proliferation in a systemic environment that does not rely on Insulin signaling? In this environment proliferating cells must solve two problems: (1) They must obtain and take up the right nutrients and (2) they must drive protein translation and growth under Insulin-limiting conditions. To address the first question, we investigated if *egr-*driven Insulin restriction may induce nutrient release from the fat body, the largest nutrient storage organ in larvae. Previous studies have shown that fat body breakdown provides nutrients for tumor growth, thereby revealing an ancient program of nutrient supply under inflammatory conditions (Ding et al, 2021; Figueroa-Clarevega & Bilder, 2015; Katheder & Rusten, 2017; Khezri et al, 2021; Lodge et al, 2021; Newton et al, 2020; Song et al, 2019). To understand if similar catabolic changes may be induced in *egr-*expressing larvae, we employed Nile Red staining to visualize lipid droplets within the fat body. We observed morphological changes, such as increased droplet size and ‘roundness’ (defined as the relationship between area and length of major axis), consistent with molecular changes associated with lipid mobilization (**Fig 3A-C, Fig S3.1 A**) (Beller et al, 2010; Figueroa-Clarevega & Bilder, 2015; Gutierrez et al, 2007; Lodge et al, 2021; Ugrankar et al, 2019). Correspondingly, fat bodies of these larvae exhibited a significant decrease in triglyceride content, similarly to fat bodies from starved larvae (**Fig 3D,E**). Moreover, we find that levels of total and activated p-Akt in fat body from *egr-*expressing larvae were reduced, similar to insulin restriction observed in starved larvae (**Fig S3.1B**). Additionally, levels of nuclear dFOXO-GFP were elevated (**Fig. S3.1 C-E**), supporting the notion that *egr-*induced insulin restriction reduces systemic Insulin signaling and promotes a catabolic fat body state.

**Figure 3.**
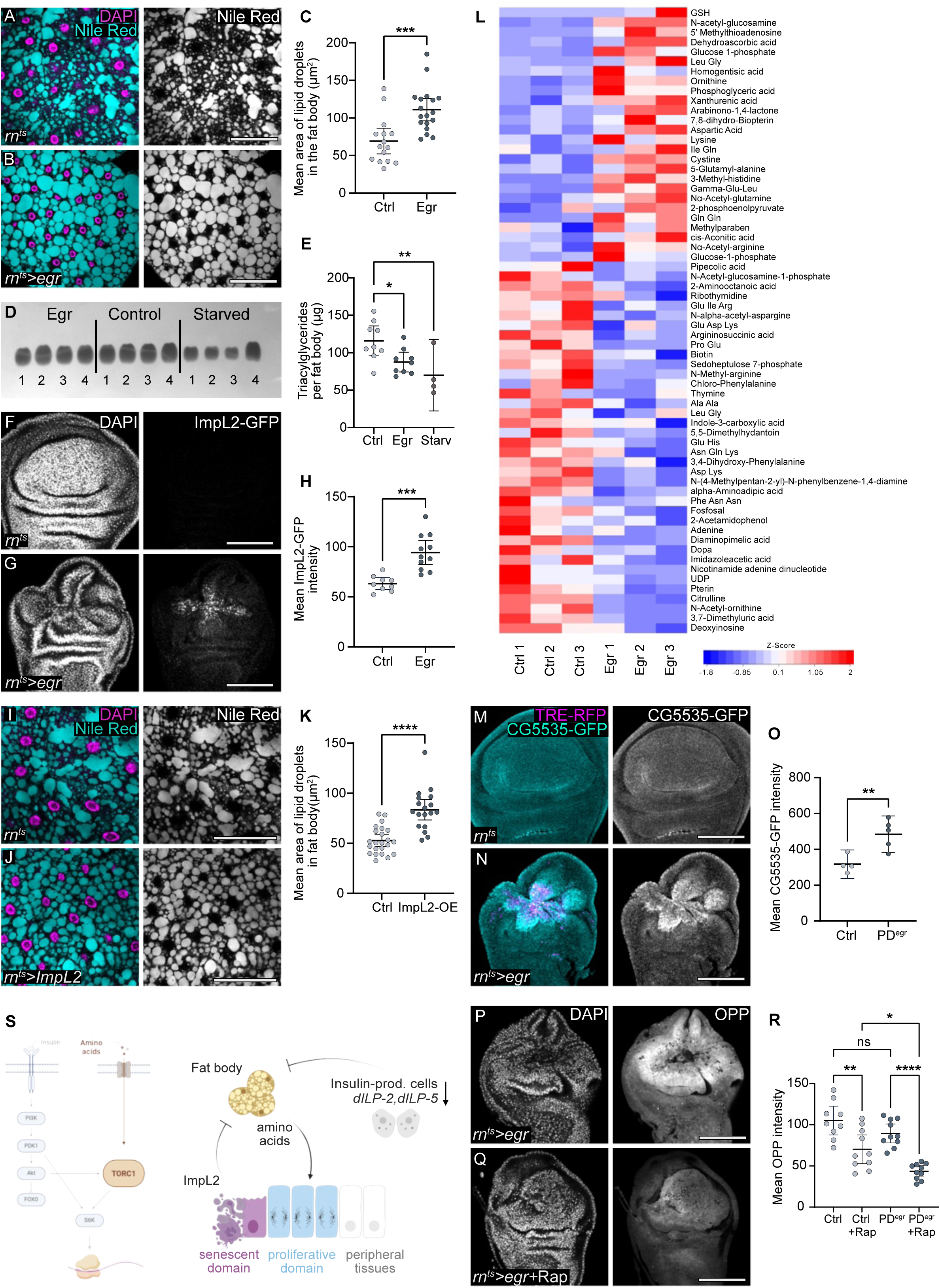
Fat body catabolism and nutrient release to hemolymph facilitates regenerative proliferation through mTORC signaling pathway. **A, B.** Nile Red staining (cyan or grey) of fat body, dissected from larvae with control (A) or *egr-*expressing (B) wing discs. Fat bodies were stained with DAPI (magenta) to visualize nuclei. **C.** Quantification of area of lipid droplets in the fat body dissected from larvae with control (A) or *egr-*expressing (B) wing imaginal discs. Mean and 95% CI are shown. Statistical significance of was tested using a two-tailed Unpaired t-test. Mean area, p-value = 0.0004. (control: n=15, *egr-*expression in discs: n=18). **D, E.** Triacylglycerides (TAG) levels were measured in *Drosophila* fat bodies dissected from third-instar larvae with *egr-*expressing wing disc, control larvae or starved larvae using thin-layer chromatography (TLC). Fat bodies from third instar larvae starved for 16hrs were used as a positive control. TAG levels were quantified per fat body in control, *egr-*expressing and starved larvae. Mean and 95% CI are shown. Statistical significance was tested using a one-way ANOVA followed by Dunnett’s test for multiple comparison.(control larvae: n=9, *egr-*expressing larvae: n=9 and starved larvae: n=4). p-values: control vs egr = 0.0353; control vs starved =0.0076. **F, G.** Expression of ImpL2-GFP in control (A) and *egr-*expressing discs (B). Discs were stained with DAPI to visualize nuclei. **H**. Quantification of ImpL2-GFP intensity in control (**F**) and *egr-*expressing discs (**G**). Mean and 95% CI are shown, and statistical significance was tested using two-tailed Welch’s t-test, p-value = 0.0001 (control: n=9, *egr-*expressing discs: n=11) **I, J.** Nile Red staining of fat body (cyan or grey), dissected from larvae with control (I) or ImpL2-expressing under *rn-*GAL4 (J) in wing imaginal discs. Fat bodies were stained with DAPI (magenta) to visualize nuclei. **K.** Quantification of area of lipid droplets in the fat body dissected from larvae with either control discs or wing imaginal discs expressing ImpL2 for 24 h under the control of *rn*-GAL4. For lipid droplet area, statistical significance was tested using the two-tailed Mann-Whitney test, p-value < 0.0001 (control: n=23, experiment: n=18). Mean and 95% CI are shown. **L.** Heat map representing relative concentration changes of metabolites identified in the larval hemolymph isolated from control larvae and *egr-*expressing larvae. The heat map was generated using quantile normalized data, and statistical analysis was performed using a two-sided, unpaired Wilcoxon rank-sum test (see Materials and Methods) and metabolites with at least <0.75 and >1.5-fold change were selected for further analysis. Metabolites in the heat map are ordered based on the log2 fold change between control and *egr-*expressing larval samples and Z-scores were visualized. Sample size was n=3 for both control and *egr-*expressing larvae. **M, N.** Expression of CG5535-GFP in control (cyan or grey) (M) and *egr-*expressing discs (N). TRE-RFP visualizes JNK-pathway activity (magenta). **O.** Quantification of mean CG5535-GFP intensity quantified in the pouch of control discs and the proliferative domain of *egr-*expressing discs (PD^egr^). Mean and 95% CI are shown. Statistical significance was tested using the two-tailed Unpaired t-test, p-value = 0.0092 (control: n=4, *egr-*expressing discs: n=5). **P, Q.** Protein synthesis visualized by OPP incorporation in *egr-*expressing wing discs from larvae fed on food without (P) or with rapamycin (200 μM) (Q) for 24 h during the 24 h period of *egr-*expression in wing discs. **R.** Quantification of mean OPP intensity in the pouch of control discs and the proliferative domain of *egr-*expressing discs (PD^egr^) from larvae fed on food without or with rapamycin (200 μM) for 24 hours during the 24 h period of *egr-*expression in wing discs. Mean and 95% CI are shown. Statistical significance was tested using a one-way ANOVA followed by Tukey’s post-hoc test for multiple comparisons (control: non-fed: n=9, fed: n=10; *egr-*expressing disc: non-fed: n=10, fed: n=10). PD^egr^ vs PD^egr^ + Rapamycin: p < 0.0001, Control + Rapamycin vs PD^egr^ + Rapamycin: p = 0.0167 and Control vs Control + Rapamycin: p = 0.0017. **S**. A scheme highlighting the branch of TORC1 signaling driving S6K activation. Model illustrates how ImpL2 is secreted from senescent-like cells in the high JNK signaling domain, repressing fat body anabolism possibly via inducing insulin resistance, and thereby inducing nutrient mobilization from fat body. This branch may reinforce the metabolic switch induced by Insulin restriction. The released nutrients such as amino acids, enter the hemolymph and are subsequently taken up by the proliferative domain cells promoting mTORC1 growth signaling and S6K activation. Scale bars: 100 μm. Fluorescence intensities are reported as arbitrary units.

Notably, JNK-signaling cells in *egr-*expressing discs also express *ImpL2*, *upd2* and *upd3*, factors previously implicated in fat body break-down during tumor growth in larval and adult hosts (**Fig 3 F-H, Fig S3.1 F-H, Fig S3.2**) (Ding et al, 2021; Figueroa-Clarevega & Bilder, 2015; Kwon et al, 2015; Romao et al, 2021). When we ectopically expressed ImpL2 for 24 h using the wing-pouch-specific driver *rn-GAL4*, we found that this was sufficient to induce lipid droplet changes consistent with fat body catabolism (**Fig 3I-K, Fig S3.1 I**). Moreover, *egr*-expressing larvae display activation of JAK/STAT signaling in fat body, which has also been associated with fat body catabolism (**Fig S3.1 J-L**) (Ding et al, 2021; Hersperger et al, 2024; Shin et al, 2020). Of note, we did not observe alterations in muscle morphology or evidence of autophagy in imaginal discs, suggesting that these tissues do not serve as a primary source for nutrients after just 24 h of *egr*-expression, different to what is observed in the chronic presence of tumors (**Fig S3.1 M-P**) (Ding et al, 2021; Figueroa-Clarevega & Bilder, 2015; Katheder & Rusten, 2017; Khezri et al, 2021; Lodge et al, 2021; Newton et al, 2020; Song et al, 2019). Combined, we conclude that the combinatorial expression of systemically acting cytokines from JNK-signaling cells in *egr-*expressing discs drives detectable nutrient mobilization from the fat body via multiple pathways.

To determine if the observed break-down of the fat body could also generate amino acid compounds supporting the high rates of protein translation in the proliferative domain, we performed an untargeted metabolomic analysis of the hemolymph from control and *egr-*expressing larvae. Our results revealed a a shift in the composition of amino acids in in *egr-*expressing larval hemolymph. We observed an enrichment of many amino acids including Leucine, Arginine or Glutamine, which are reported to activate mTORC1 (Lama-Sherpa et al, 2023; Yue et al, 2022). Surprisingly, many come enriched in form of mixed dipeptide species. Furthermore, elevated levels of dipeptide species and free amino acids associated with glutamate metabolism were observed, including increased concentrations of glutamine, glutamate, and glutamyl dipeptides (**Fig 3 L**). Notably, a signature of arginine metabolism emerged, characterized by increased levels of ornithine, N-acetyl-arginine, and aspartate alongside decreased levels of argininosuccinic acid, citrulline, and N-acetyl-ornithine. This altered metabolite profile in the hemolymph represents a flexible supply of building blocks not only for protein synthesis, but also for energy production, co-factor generation, and glutathione-based redox metabolism (Altman et al, 2016; Keshet et al, 2018; Lieu et al, 2020; Marti & Reith, 2021).

We next examined the ability of proliferating cells to absorb these amino acids. We investigated the expression of solute carrier (SLC) transporters and found that several transcripts were expressed and elevated in *egr-*expressing discs. For instance, *CG15279* (a putative glycine and proline transporter from the SLC6 family), *path* (a potential alanine and glycine transporter from the SLC36 family), *mnd* (an amino acid/polyamine transporter involved in leucine import), and *CG5535* (a putative lysine, arginine, and ornithine transporter from the SLC7 family) were all elevated (**Fig S3.2**). Upregulation of CG5535 was confirmed by immunofluorescence, and we similarly detected increased levels of CG1139 (Arcus), another SLC36 family member that may transport alanine, glycine, and proline (**Fig 3M-O, Fig S3.1 Q,R**). In addition, a potential sugar transporter, *CG3168* of the SLC22 family, and TRET-1, which is predicted to import trehalose, were strongly upregulated (**Fig S3.1 S-U**) These findings suggest that proliferating cells have an enhanced capacity for importing both amino acids and energy sources.

One regulatory branch promoting protein translation in response to amino acid availability is mediated by mTORC1 (Jewell et al, 2013; Lama-Sherpa et al, 2023; Liu & Sabatini, 2020; Shimobayashi & Hall, 2016; Yue et al, 2022). Moreover, mTORC1 activity itself is promoted by sugar import, boosting ATP production, and hence inhibiting the mTORC1 antagonist AMPK (Jewell et al, 2013; Liu & Sabatini, 2020; Shimobayashi & Hall, 2016). The observed upregulation of amino acid and sugar transporters in *egr-*expressing discs could therefore facilitate mTORC1 activation. To demonstrate that the mTORC1 pathway supports protein synthesis during wing disc regeneration, we inhibited mTORC1 activity by feeding *egr-*expressing larvae the mTOR inhibitor Rapamycin for 24 h during *egr-*expression (Li et al, 2014; Wu et al, 2022). This treatment resulted in a pronounced reduction in protein synthesis within the proliferative domain of *egr-*expressing discs, demonstrating that mTORC1 activity is essential for protein translation during regeneration, as in wild type control discs (**Fig 3P-R, Fig S3.1 V,W**). Overall, our data support a model in which fat body catabolism, enhanced amino acid uptake and mTORC1 activation together support the rapid growth and proliferation of the regenerating domain (**Fig 3S**).

### (4) Pdk1 is upregulated in the proliferating domain

S6K, an important effector of mTORC1, directly activates protein translation (Liu & Sabatini, 2020; Wu et al, 2022). S6K transcripts are upregulated in the wound-associated cell populations of *egr-*expressing discs (**Fig S3.2D**) (Floc’hlay et al, 2023), suggesting that S6K is positively regulated during tissue repair. Importantly, staining for phosphorylated ribosomal S6 (p-S6), a direct target of S6K, revealed that the proliferating domain of *egr-*expressing discs exhibited strongly elevated p-S6 levels (**Fig 4A-C**). However, elevated p-S6 levels cannot be explained solely by mTORC1 activation of S6K because S6K requires co-activation by the Pdk1, a kinase central to canonical Insulin/PI3K signaling (Hopkins et al, 2020; Hoxhaj & Manning, 2020; Wu et al, 2022).

**Figure 4.**
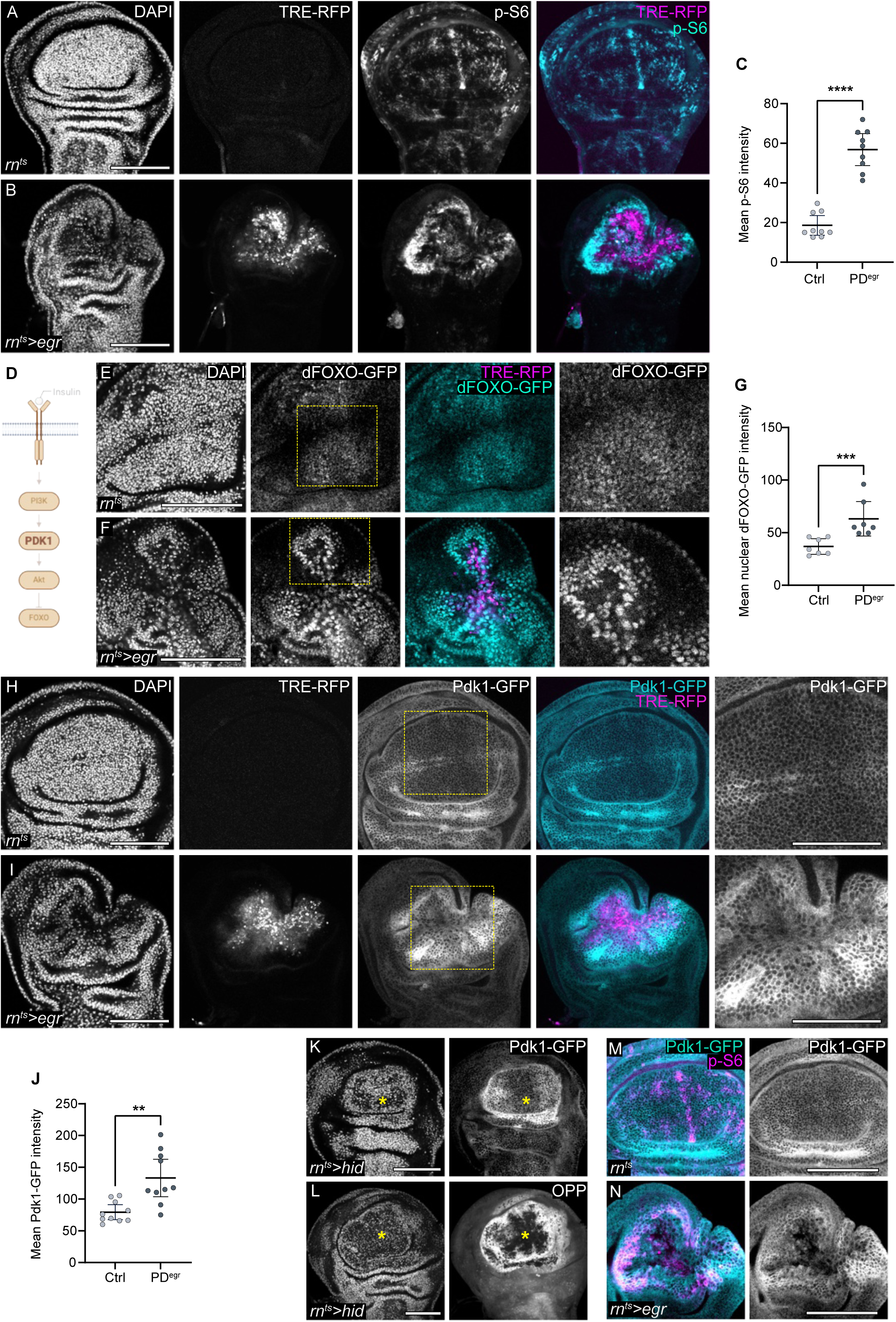
Pdk1 is upregulated in the proliferative domain. **A, B.** Immunostaining for phosphorylated-S6 (pS6, cyan or grey) in control (A) and *egr-*expressing wing discs (B). TRE-RFP visualizes JNK-pathway activity (magenta or grey). Discs were stained with DAPI to visualize nuclei. **C**. Quantification of mean p-S6 intensity in the pouch of control discs and the proliferative domain of *egr-*expressing discs (PD^egr^). Mean and 95% CI are shown. Statistical significance was tested using the two-tailed Mann-Whitney test, p-value < 0.0001 (control: n=9, *egr-*expressing discs: n=9). **D**. Schematic representation of canonical insulin signaling pathway, highlighting the insulin/Akt/FOXO axis. Phosphatidylinositol kinase-1 (Pdk1) is key serine/threonine kinase in this pathway. **E, F.** Expression of dFOXO-GFP (cyan or grey) in control (E) and *egr-*expressing wing discs (F). TRE-RFP visualizes JNK-pathway activity (magenta or grey), and discs were stained with DAPI to visualize nuclei. Yellow dotted box marks the inset region. BDSC: 38644 fly line was used. **G.** Quantification of mean nuclear dFOXO-GFP intensity in the pouch of control discs or the proliferative domain of *egr-*expressing discs (PD^egr^). Mean and 95% CI are shown, and statistical significance was tested using the two-tailed Mann-Whitney test, p-value = 0.0006 (control: n=7, *egr-*expressing disc: n=7). **H, I.** Expression of Pdk1-GFP (cyan or grey) in control (H) and *egr-*expressing discs (I). TRE-RFP visualizes JNK-pathway activity (magenta or grey), and discs were stained with DAPI to visualize nuclei. Yellow dotted box marks the inset region. **J.** Quantification of Pdk1-GFP intensity in the pouch of control discs or the proliferative domain of *egr-*expressing discs (PD^egr^). Statistical significance was tested using a two-tailed Welch’s t-test, p-value = 0.0025 (control: n=10, *egr-*expressing disc: n=10). Mean and 95% CI are shown. **K.** Expression of Pdk1-GFP in pro-apoptotic gene *hid-*expressing discs and DAPI visualizes nuclei. The yellow asterisk marks the damage or *hid*-expressing region. Discs were stained with DAPI to visualize nuclei. **L.** Protein synthesis visualized by OPP incorporation in *hid-*expressing discs and DAPI visualizes nuclei. The yellow asterisk marks the damage or *hid*-expressing region. Discs were stained with DAPI to visualize nuclei. **M, N.** Expression of Pdk1-GFP (cyan or grey) and p-S6 (magenta) in control (M) and *egr-*expressing wing discs (N). Scale bars: 100 μm. Fluorescence intensities are reported as arbitrary units.

However, how can proliferating cells promote activation of S6K by Pdk1, considering that *egr-*expressing larvae exhibit insulin restriction? To understand how Pdk1 may be activated, we examined if canonical Insulin/PI3K signaling was active in the proliferating domain of *egr-*expressing discs. We therefore visualized the nuclear localization of dFOXO, the most downstream effector of Insulin/PI3K activity, using two independent GFP-tagged lines. We observed high nuclear levels of dFOXO in the proliferating domain of *egr-*expressing discs, suggesting that Insulin/PI3K signaling there is low, which is consistent with our evidence for systemic Insulin restriction (**Fig 4 D-G, Fig S4A-C**). This conclusion is further supported by unchanged InR levels and even reduced levels of Akt and phosphorylated Akt (P-Akt) in the proliferative domain (**Fig S4D-K**). In contrast, we found that Pdk1 expression is strongly upregulated in the proliferative domain (**Fig 4H-J**) (see Materials & Methods, section *Drosophila* genetics). Similar upregulation of Pdk1 was also observed in other models of tissue damage and regeneration, including those using expression of *hid* or *reaper* to drive cell death. In these models, elevated Pdk1 levels also correlated with high protein translation, demarcating the proliferating domains (**Fig 4K,L, Fig S4L-Q**). In fact, we found that high levels of Pdk1 correlated with high levels of p-S6 staining in *egr-*expressing discs (**Fig 4M,N**). Collectively, these observations suggest that Pdk1 levels may play a central role in regenerative proliferation.

### (5) Pdk1 upregulation is sufficient to drive growth and is necessary for regenerative proliferation

To understand if Pdk1 upregulation is a central mechanism driving regenerative proliferation, we expressed either a wild-type or a constitutively active form of Pdk1 in the wing pouch for 24 hours using *rn-GAL4*. Compared to control discs, both forms of Pdk1 resulted in higher levels of protein translation and larger pouch sizes. Larger pouch sizes correlated with more cell proliferation, as EdU incorporation in Pdk1 expressing domains was elevated (**Fig 5 A-F, Fig S5A-D**). These results demonstrate that high Pdk1 levels are sufficient to support a metabolic program characteristic of regenerative proliferation. Notably, the fact that overexpression of a wild-type Pdk1 alone enhances protein synthesis and proliferation, suggests that Pdk1 activity scales with expression levels, functioning independently of other canonical upstream signaling inputs, such as Insulin/PI3K. Of note, Pdk1 can auto-activate via transphosphorylation when recruited to the plasma membrane (Komander et al, 2004; Levina et al, 2022), reflecting potentially Insulin-independent means for Pdk1activation.

**Figure 5.**
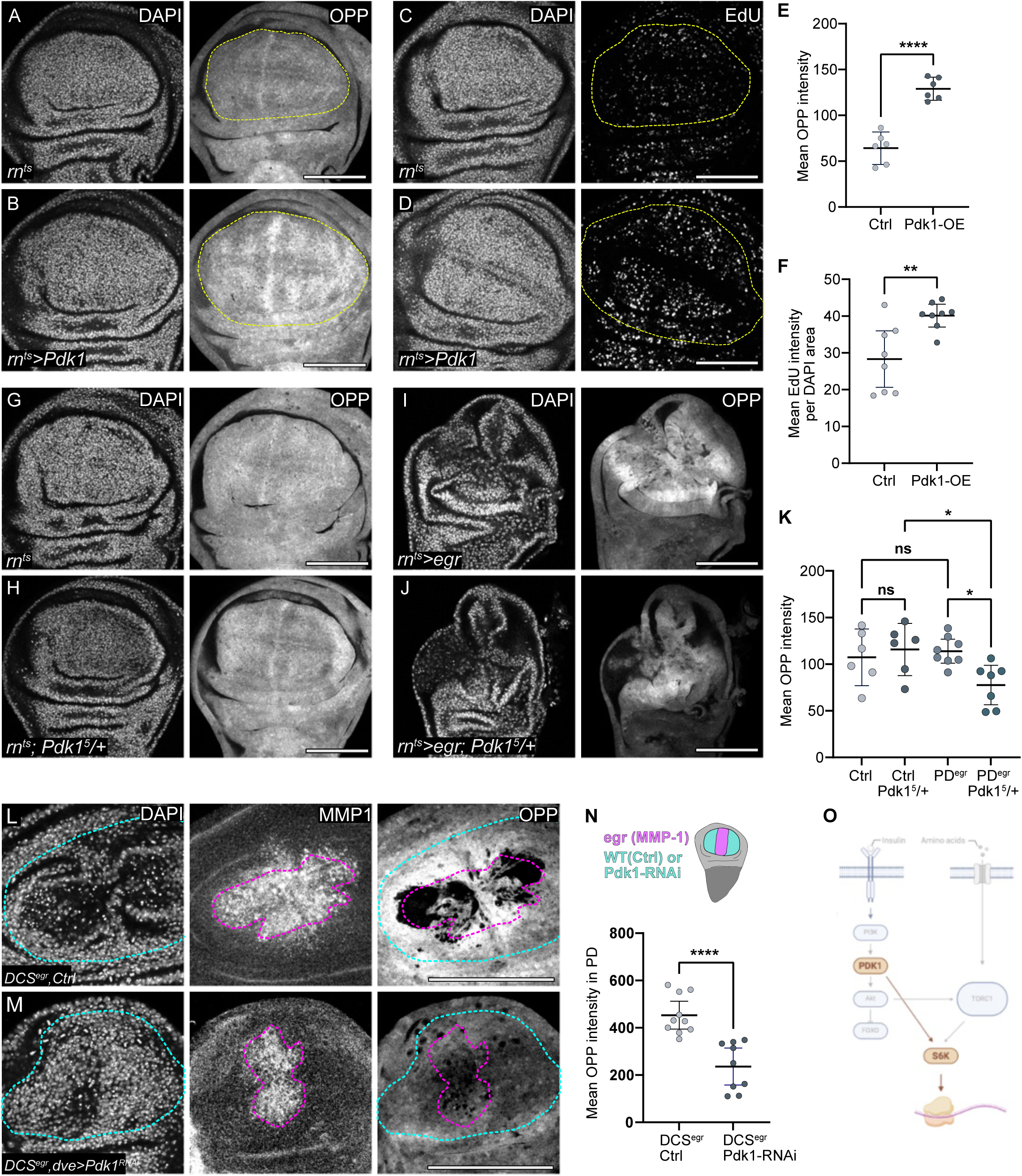
Pdk1 upregulation is sufficient and necessary for regenerative growth and proliferation. **A,B.** Protein synthesis visualized by OPP incorporation in control wing discs (A) and discs expressing Pdk1 for 24 h under the control of *rn*-GAL4 driver (B). DAPI staining visualizes nuclei. The dotted yellow line indicates the boundary of the wing pouch based on landmark tissue folds, approximating the *rn*-GAL4 expression domain **C, D.** EdU incorporation in control (C) and discs expressing UAS-Pdk1 for 24 h under the control of *rn-*GAL4 driver (D). DAPI staining visualizes nuclei. The dotted yellow line indicates the boundary of the wing pouch based on landmark tissue folds, approximating the *rn-*GAL4 expression domain **E.** Quantification of mean OPP intensity in the pouch of control and *Pdk1*-expressing wing discs. Mean and 95% CI are shown, and statistical significance was tested using a two-tailed Unpaired t-test, p-value < 0.0001 (control: n=6, *Pdk1-*expressing disc: n=6). **F.** Quantification of mean EdU intensity per DAPI area serving as a proxy for DNA replication speed, in the pouch of control and *Pdk1*-expressing wing discs. Mean and 95% CI are shown. Statistical significance was tested using the two-tailed Welch’s t-test, p-value = 0.0078 (control: n=8, *Pdk1-*expressing disc: n=8). **G-J.** Protein synthesis visualized by OPP incorporation in control (G, H) and *egr-*expressing wing discs (I, J) that were either wild type (G, I) or heterozygous mutant for the *Pdk1^5^* null allele (H, J). Discs were stained with DAPI to visualize nuclei. **K.** Quantification of mean OPP intensity in the pouch of control or the proliferative domain of *egr-*expressing discs (PD^egr^) that were either wild type or heterozygous mutant for the *Pdk1^5^* null allele. Statistical significance was tested using a one-way ANOVA followed by Tukey’s post-hoc test for multiple comparisons. (control:n=6, control, *Pdk1^5^*/+: n=6; *egr*:n=8, *egr, Pdk1^5^/+*: n=7). PD^egr^ vs PD^egr^*: Pdk1^5^*/+: p-value = 0.0300 and Control: *Pdk1^5^*/+ vs PD^egr^ *: Pdk1^5^*/+: p-value= 0.0350. Mean and 95% CI are shown. **L, M.** DUAL Control genetic ablation system (DCS) was used to manipulate gene expression in the proliferative domain. Early third instar (L3) larvae were heat shocked at 37°C for 1h and dissected after 24 h. A single heat shock activates both ablation (region inside the magenta dotted line) in the *salm* domain and gene manipulation in the proliferative domain via *dve-*GAL4 (the *dve-GAL4* expressing region is marked with a cyan dotted line). MMP1 is a target gene of JNK activated by eiger expression, thereby demarcating the ablated domain. Protein synthesis is visualized by OPP incorporation in wing imaginal discs in control (L) and Pdk1 knockdown (M) wing imaginal disc. **N.** Quantification of mean OPP intensity in the proliferative domain using a 20 μm band around the ablated region of control (*DCS^egr^*, ctrl) and *Pdk1* knockdown (*DCS^egr^, Pdk1 RNAi*) wing imaginal discs with *egr* expression in the *salm* domain. On top, a simplified schematic illustrating the *egr-*expressed region (magenta) and the *dve-GAL4* expressing region (cyan) used for Pdk1 knockdown. Statistical significance was assessed using a two-tailed unpaired *t*-test (*p* < 0.0001). (*DCSegr*, control: *n* = 10; *DCSegr, Pdk1 RNAi*: *n* = 9). Mean and 95% CI are shown. **O.** A scheme highlighting the branch of Pdk1 signaling driving S6K activation. Scale bars: 100 μm. Fluorescence intensities are reported as arbitrary units.

To provide evidence that Pdk1 function is also necessary to support proliferation in *egr-*expressing discs, we genetically reduced Pdk1 function by establishing heterozygosity for a null allele of *Pdk1* (Rintelen et al, 2001). In control discs, this heterozygosity did not affect overall protein synthesis compared to wild type discs, indicating that under normal developmental conditions a single copy of *Pdk1* is sufficient for growth. However, protein synthesis in the proliferating domain of *egr-*expressing discs heterozygous for *Pdk1* was significantly reduced (**Fig 5G-K**). This finding demonstrates that the proliferating domain is particularly sensitive to reductions in Pdk1 levels and underscores the necessity for elevated Pdk1 to support protein synthesis during regeneration. To provide further proof, we expressed a validated RNAi against *Pdk1* specifically in proliferating cells of *egr*-expressing discs under control of the DUAL Control genetic ablation system (**Fig S5E,F**) (Harris et al, 2020). Consistent with our heterozygous mutant results, *Pdk1* knockdown led to a marked decrease in protein translation in the proliferating domain, confirming a cell type specific function for Pdk1 upregulation in the proliferating domain during regeneration (**Fig 5L-N**). Overall, these findings establish that Pdk1 upregulation is not only sufficient to drive protein translation and growth but is also critically required to sustain protein translation in the proliferative domain of regenerating imaginal discs under conditions of systemic insulin restriction (**Fig 5O**).

### (6) Pdk1 is regulated by JAK/STAT signaling

To understand how Pdk1 levels are controlled during regeneration, we examined the role of JAK/STAT signaling, which is activated in the proliferating domain of *egr-*expressing discs to promote cell proliferation and survival (**Fig 6A,B**) (Cosolo et al, 2019; Floc’hlay et al, 2023; Harris et al, 2020; Jaiswal et al, 2023; Worley et al, 2018; Worley et al, 2022). We therefore expressed STAT92E in the posterior compartment of wing imaginal discs, which elevates JAK/STAT signaling, using the *en-GAL4* driver (Jaiswal et al, 2023). This manipulation caused an increase in levels of Pdk1-GFP, phospho-S6 and OPP incorporation (**Fig 6C-H, Fig S6C-E**), demonstrating that JAK/STAT activation is sufficient to promote Pdk1 upregulation, S6 activation and protein translation. Conversely, when we expressed a RNAi construct to knock down *STAT92E* in the posterior compartment during normal development, we observed a decrease in both Pdk1-GFP levels and OPP incorporation, suggesting that JAK/STAT activity can limit Pdk1 levels and protein translation (**Fig 6 I-L**). Importantly, the JAK/STAT-Pdk1-S6 axis is also essential for regeneration: we genetically reduced *STAT92E* function in *egr-*expressing discs by establishing heterozygosity for a null allele of *STAT92E* and found that it reduced OPP incorporation specifically in *egr*-expressing discs but not in control discs (**Fig 6M-O, Fig S6F,G**). Moreover, when we expressed a RNAi against *STAT92E* specifically in proliferating cells of *egr*-expressing discs under control of the DUAL Control genetic ablation system (Harris et al, 2020), the elevated protein translation was abrogated (**Fig 6P-R, Fig S6H,I**). Combined, these findings reveal a novel function for JAK/STAT signaling in tissue repair and regeneration by promoting Pdk1 upregulation which, together with mTORC1, supports insulin-independent growth in the proliferating domain (**Fig S6J**).

**Figure 6.**
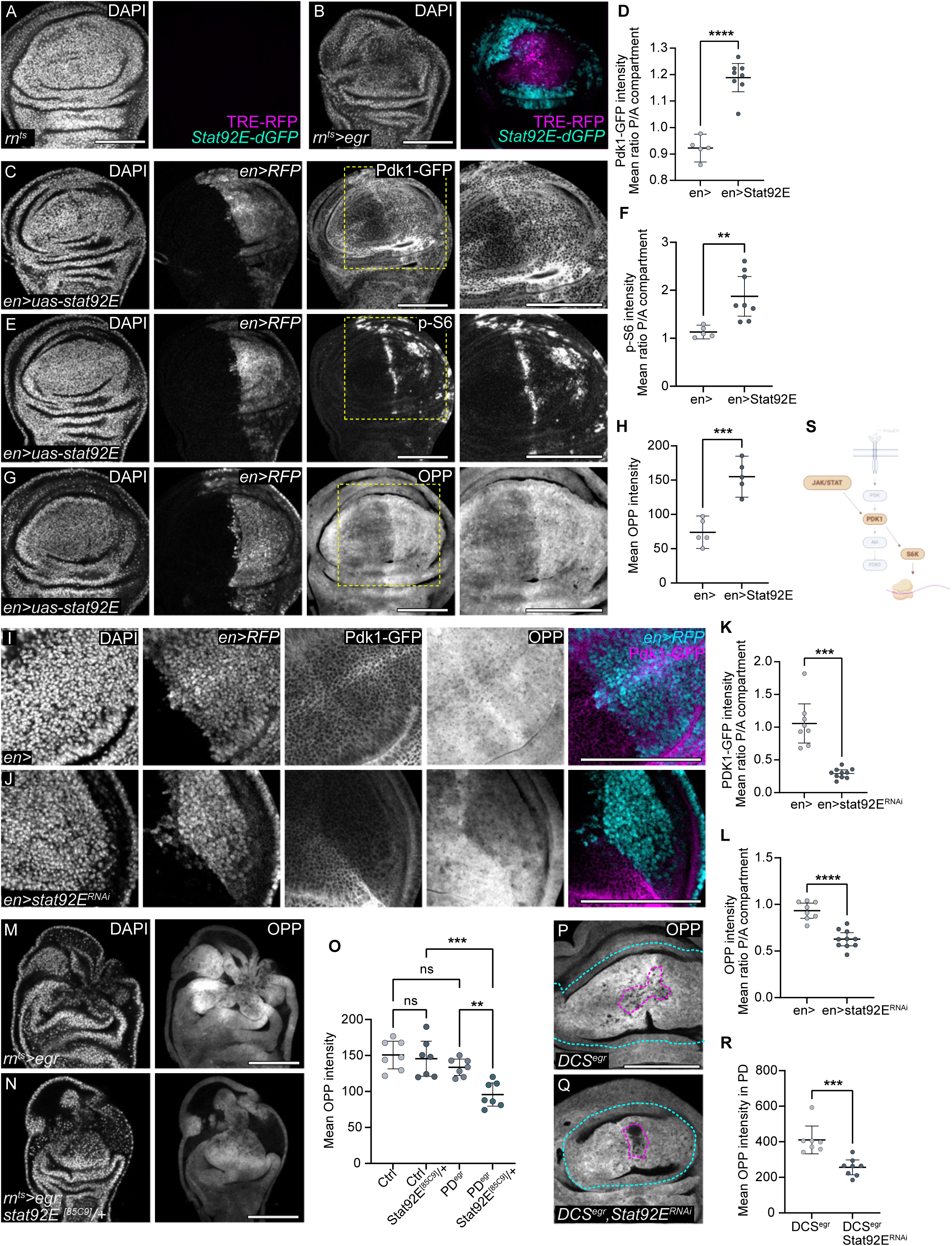
Pdk1 is regulated by JAK/STAT signaling. **A, B.** A control and *egr-*expressing wing imaginal disc with TRE-RFP (magenta) and Stat92E-dGFP (cyan) reporter. Discs were stained with DAPI to visualize nuclei. **C, E, G.** Wing imaginal discs expressing *STAT92E* in the posterior compartment under the control *en-*GAL4 driver which also drives expression of UAS-RFP. Discs were assayed for expression of Pdk1-GFP (C), levels of p-S6 (E), and protein synthesis using OPP incorporation assays (G). Discs were stained with DAPI to visualize nuclei. **D, F, H.** Quantifications of levels of Pdk1-GFP (D), p-S6 (F) and protein synthesis using OPP incorporation assays (H) using the posterior to anterior (P/A) ratio in control wing imaginal discs and wing imaginal discs expressing *STAT92E* in the posterior compartment under the control *en-*GAL4. In the graph displaying Pdk1-GFP intensity (D), statistical significance was tested using the two-tailed Unpaired t-test, p-value < 0.0001 (control: n=5, experiment: n=8). In the graph displaying p-S6 intensity (F), statistical significance was tested using two-tailed Welch’s t-test, p-value = 0.0033 (control: n=5, experiment: n=8). In the graph displaying OPP intensity (H), statistical significance was tested using the two-tailed Unpaired t-test, p-value = 0.0004 (control: n=5, experiment: n=5). Mean and 95% CI are shown. **I, J.** Control wing disc (I) and wing imaginal discs expressing *STAT92E-RNAi* (J) in the posterior compartment under the control *en-*GAL4 driver which also drives expression of UAS-RFP. Discs were assayed for expression of Pdk1-GFP and protein synthesis using OPP incorporation assays. Discs were stained with DAPI to visualize nuclei. **K.** Quantifications of Pdk1-GFP levels using the posterior to anterior (P/A) ratio in control wing imaginal discs and wing imaginal discs expressing *STAT92E-RNAi* in the posterior compartment under the control *en-*GAL4 driver. Statistical significance was tested using the two-tailed Welch’s t-test, p-value = 0.0004 (control: n=8, experiment: n=10). Mean and 95% CI are shown. **L.** Quantifications protein synthesis levels using the OPP posterior to anterior (P/A) ratio in control wing imaginal discs and wing imaginal discs expressing *STAT92E-RNAi* in the posterior compartment under the control *en-*GAL4 driver. Statistical significance was tested using the two-tailed Unpaired t-test, p-value < 0.0001 (control: n=8, experiment: n=10). Mean and 95% CI are shown. **M, N.** Protein synthesis visualized by OPP incorporation in *egr-*expressing discs that were either wild type (M) or heterozygous mutant for the *Stat92E^85C9^*null allele (N). Discs were stained with DAPI to visualize nuclei. **O.** Quantification of mean OPP intensity in the pouch of control and the proliferative domain of *egr-*expressing discs (PD^egr^) that were either wild type or heterozygous mutant for the *Stat92E^85C9^* null allele. Statistical significance was tested using a one-way ANOVA followed by Tukey’s post-hoc test for multiple comparisons. (control:n=7, control, *Stat92E^85C9^*/+: n=7; *egr*:n=7, *egr, Stat92E^85C9^/+*: n=7). PD^egr^ vs PD^egr^ *:Stat92E^85C9^*/+: p-value = 0.0080 and Control*: Stat92E^85C9^*/+ vs PD^egr^ *: Stat92E^85C9^*/+: p-value =0.0005. Mean and 95% CI are shown. **P, Q.** A DUAL Control genetic ablation system (DCS) was used to manipulate gene expression in the proliferative domain. Early third instar (L3) larvae were heat shocked at 37°C for 75min and dissected after 40 h. A single heat shock activated both ablation (region inside the magenta dotted line) in the *salm* domain and gene manipulation in the proliferative domain via *dve-*GAL4 (*dve-GAL4* expressing region is marked with a cyan dotted line). Protein synthesis visualized by OPP incorporation in control (P) and *Stat92E* knockdown (Q) wing imaginal disc. **R.** Quantification of mean OPP intensity in the proliferative domain using a 20 μm band around the ablated region of control (*DCS^egr^*) and *STAT92E* knockdown (*DCS^egr^, Stat92E-RNAi*) wing imaginal discs with *egr* expression in the *salm* domain. Mean and 95% CI are shown. Statistical significance was assessed using a two-tailed Mann-Whitney’s test, *p value* = 0.0006. (*DCSegr*, control: *n* = 7; *DCSegr, Stat92E-RNAi*: *n* = 8). **S.** A scheme highlighting that JAK/STAT activation positively affects Pdk1 activity towards S6K. Scale bars: 100 μm. Fluorescence intensities are reported as arbitrary units.

### (7) Pdk1 upregulation, protein translation and JAK/STAT signaling are linked in tumor growth

We wanted to understand, if the JAK/STAT-Pdk1-S6 signaling axis observed during regenerative proliferation also plays a role in tumor growth, given that tumors often coopt physiological repair processes for their pathological purposes. We thus analyzed imaginal discs in which tumorigenesis was induced by both reducing the function of the tumor-suppressor gene *scrib* (Bilder et al, 2000) and ectopically expressing oncogenic *Ras^V12^* (Dillard et al, 2021; Pagliarini & Xu, 2003). We used the *rn-GAL4* driver to drive expression of *scrib-RNAi* and *Ras^V12^* in the entire wing pouch, which causes pronounced overproliferation (Cong et al, 2021; Jaiswal et al, 2023; Khezri et al, 2021). This tumor model is characterized by the coordinated but spatially separated activation of JNK and JAK/STAT signaling networks, whose regulatory and functional characteristics resemble those active during tissue repair in *egr-*expressing discs (**Fig S7A**) (Jaiswal et al, 2023). In *Ras^V12^,scrib-RNAi* expressing discs, we found striking correlation between regions of JAK/STAT activation, elevated protein translation and increased Pdk1 level (**Fig 7A-F**). An independent tumor model of *Psc-Su(z)2* tumors also shows correlated elevation of JAK/STAT activation and protein synthesis (**Fig S7B,C**). Importantly, these tumors, like *egr-*expressing discs, induce all hallmarks of systemic cachexia, including reduced protein translation in the non-transformed notum (**Fig 7G-I**), and must therefore overcome growth restrictions imposed on peripheral tissues (Hodgson et al, 2021; Katheder & Rusten, 2017; Khezri et al, 2021). Our observation demonstrate that tumors likely coopt JAK/STAT activation and Pdk1 elevation to support protein translation and thus tumor proliferation.

**Figure 7.**
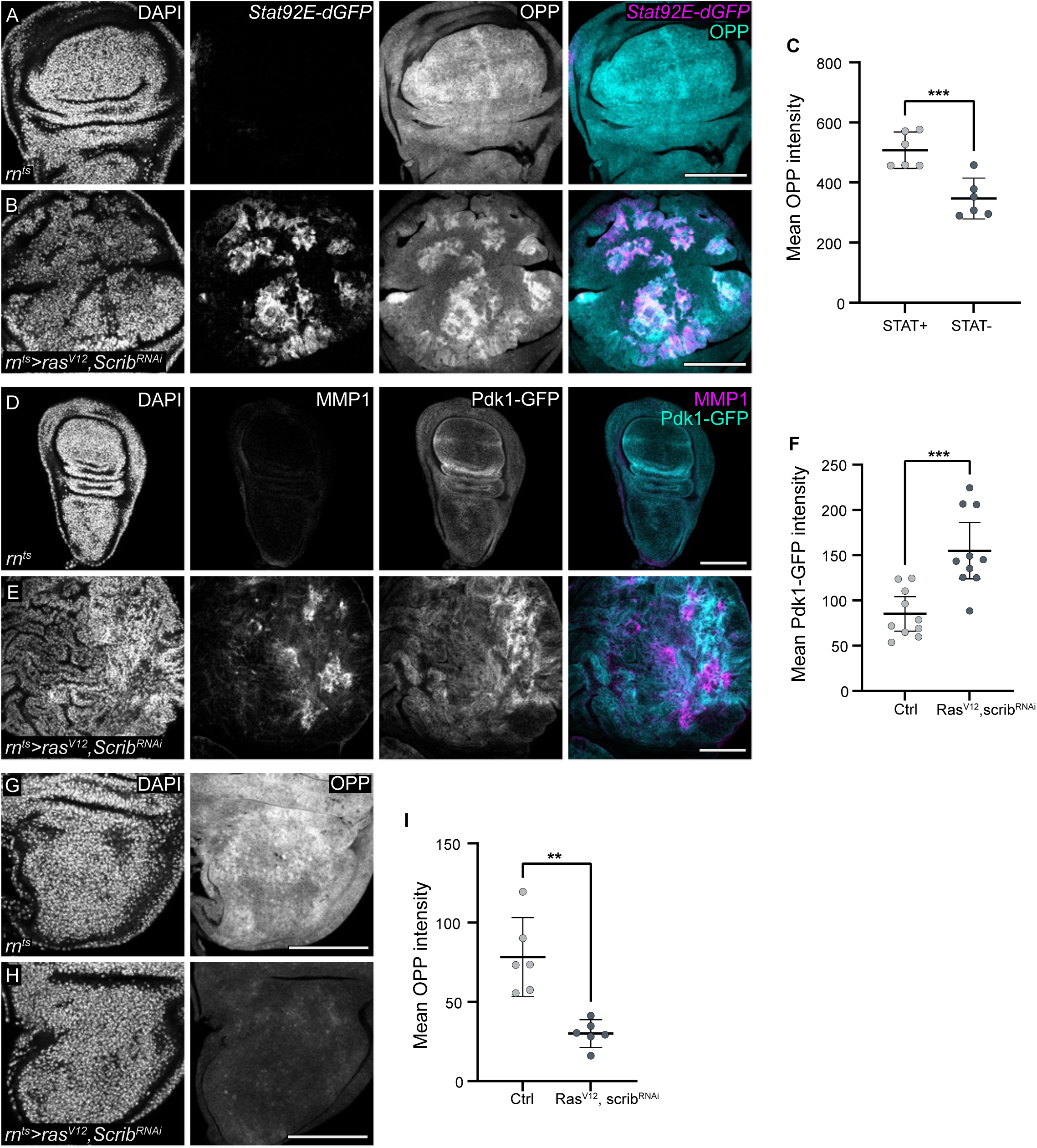
Pdk1 upregulation, protein translation and JAK/STAT signaling are linked in tumor growth. **A, B.** Control wing disc (A) and wing disc expressing *Ras^V12^, scrib-RNAi* (B) for 44h starting at Day6 AED. Protein synthesis is visualized by OPP incorporation (cyan or grey) in control (A) and *Ras^V12^, scrib-RNAi* expressing wing disc (B) along with expression of the JAK/STAT reporter Stat92E-dGFP (magenta or grey). Discs were stained with DAPI to visualize nuclei. **C**. Quantification of mean OPP intensity in the STAT positive and STAT negative region within a *Ras^V12^, scrib-RNAi* expressing wing discs. Mean and 95% CI are shown. Statistical significance was tested using the two-tailed Paired t-test, p-value = 0.0006 (control: n=6, experiment: n=6). **D, E.** Control wing disc and wing disc expressing *Ras^V12^, scrib-RNAi* for 44h starting at Day6 AED. Expression of Pdk1-GFP (cyan or grey) in control (D) and *Ras^V12^, scrib-RNAi* expressing wing disc (E) along with MMP1 staining to visualizes JNK-pathway activity (magenta or grey). Discs were stained with DAPI to visualize nuclei. **F**. Quantification of mean Pdk1-GFP intensity in the control disc pouch and *Ras^V12^, scrib-RNAi* expressing wing discs pouch. Statistical significance was tested using the two-tailed Unaired t-test, p-value = 0.0004 (control: n=10, experiment: n=10). Mean and 95% CI are shown. **G, H**. Protein synthesis visualized by OPP incorporation in the notum of control wing disc (G) and wing disc expressing *Ras^V12^, scrib-RNAi* (H) for 44h starting at Day6 AED. Discs were stained with DAPI to visualize nuclei. **I.** Quantification of mean OPP intensity in the notum of control wing disc and wing disc expressing *Ras^V12^, scrib-RNAi*. Statistical significance was tested using the two-tailed Welch’s t-test, p-value = 0.0031 (control: n=6, experiment: n=6). Mean and 95% CI are shown. Scale bars: 100 μm.

### (8) Pdk1 downregulation in peripheral discs correlates with systemic growth restriction

Strikingly, Pdk1 levels appear to be a target of systemic regulation during Insulin restriction in imaginal discs. In *egr-*expressing larvae, we consistently observed that Pdk1 levels were significantly lower in peripheral imaginal disc, including the notum, leg and eye, if compared to wild type control discs (**Fig 8A-F, Fig S8**). This observation was not limited to *egr-*expressing larvae; even in the peripheral imaginal discs of *Ras^V12^,scrib-RNAi* tumor-bearing larvae, Pdk1 levels were markedly reduced (**Fig 8G-I**). This targeted reduction of Pdk1 in peripheral imaginal discs is consistent with the overall decrease in protein synthesis and growth observed in these regions. Such downregulation suggests that systemic signals, maybe even Insulin restriction, actively suppress Pdk1 expression in non-regenerative areas. In contrast, JAK/STAT signaling in regenerative areas can counteract Pdk1 suppression and locally elevate Pdk1 levels and function. This dual regulation –upregulation in regenerative zones and downregulation in non-essential areas –highlights Pdk1’s central role in mediating the balance between local regenerative demands and systemic metabolic reprogramming. Consequently, Pdk1 emerges not only as a key driver of insulin-independent regenerative proliferation but also as an integrative node that coordinates systemic growth control during tissue repair and tumorigenesis.

**Figure 8.**
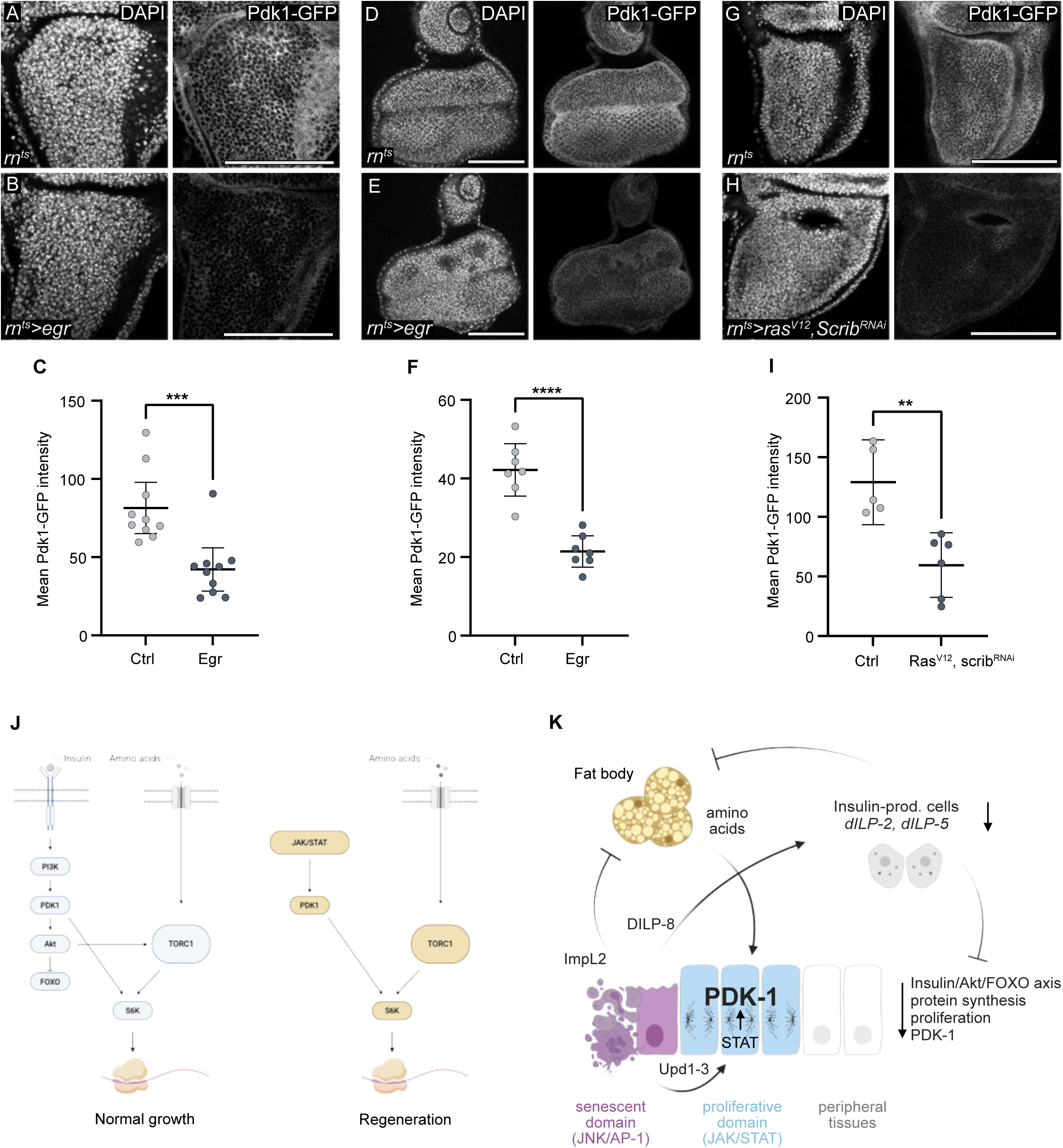
Pdk1 downregulation in peripheral discs correlates with systemic growth restriction. **A, B.** Expression of Pdk1-GFP in the notum of control (A) and *egr-*expressing discs (B). Discs were stained with DAPI to visualize nuclei. **C**. Quantification of mean Pdk1-GFP intensity in the notum of control (A) and *egr-*expressing discs (B). Mean and 95% CI are shown. Statistical significance was tested using the two-tailed Mann-Whitney’s test, p-value = 0.0007 (control: n=10, experiment: n=10). **D, E.** Expression of Pdk1-GFP in the eye imaginal disc, dissected from larvae with control (D) or *egr-*expressing (E) wing imaginal discs using *rn*-GAL4 driver. Discs were stained with DAPI to visualize nuclei. **F**. Quantification of mean Pdk1-GFP intensity in the eye imaginal disc, dissected from larvae with control (D) or *egr-*expressing (E) wing imaginal discs. Mean and 95% CI are shown. Statistical significance was tested using the two-tailed Unpaired t-test, p-value < 0.0001 (control: n=7, experiment: n=7). **G, H**. Expression of Pdk1-GFP in the notum of control wing disc (G) and wing disc expressing *Ras^V12^, scrib-RNAi* (H) for 44h starting at Day6 AED. Discs were stained with DAPI to visualize nuclei. **I.** Quantification of mean Pdk1-GFP intensity in the notum of control wing disc and wing disc expressing *Ras^V12^, scrib-RNAi*. Mean and 95% CI are shown. Statistical significance was tested using the two-tailed Unpaired t-test, p-value = 0.0022 (control: n=5, experiment: n=6). **J**. A model summary illustrating growth signaling mediated by S6K activation via canonical Insulin signaling (Insulin/AKT/FOXO and mTORC1) during normal developmental (left). On the right, growth signaling in the proliferative domain is maintained in an insulin restricted environment through the JAK/STAT-Pdk1-S6K axis, supported by mTORC1 signaling. **K.** A model summary depicting the interaction between local and systemic metabolism to support growth in the proliferative domain. Inter-organ signaling is initiated by senescent-like cells in the high JNK signaling domain at the center of tissue damage via secreted Dilp8, ImpL2 and Upd1,2,3. Dilp8 reduces insulin-like peptide expression by acting on IPCs, thereby restricting systemic insulin signaling in peripheral tissues. Additionally, ImpL2 acts on the fat body, and combined both mechanisms facilitates nutrient mobilization from fat body stores. Secreted Unpaired ligands activate JAK/STAT signaling in the nearby proliferative domain, upregulating Pdk1 and instructing S6K activation. Levels of Pdk1 in the proliferative and peripheral regions determines the tissues growth status in an insulin-restricted environment.

## Discussion

Our study reveals how signaling through a JAK/STAT-Pdk1-S6K axis promotes metabolic reprogramming to drive tissue repair and regeneration in an environment of systemic growth restriction (**Fig 8J,K**). We demonstrate that in response to inflammatory damage, a senescent subpopulation of cells at the center of tissue damage induces insulin restriction via action of dILP8 on IPCs, as well as insulin resistance via action of ImpL2 on the fat body. The resulting systemic reduction of insulin signaling is evident from the decreased expression of *dILP2* and *dILP5* in IPCs, as well as the enhanced nuclear localization of dFOXO, low rates of protein synthesis and proliferation in peripheral tissues, or reduced Akt signaling and nutrient mobilization in the fat body. These findings align with previous work suggesting that systemic growth restriction, for example via repression of Ecdysone signaling, helps synchronize regeneration with developmental timing and prevents the overgrowth of undamaged tissues (Cao et al, 2022; Colombani et al, 2012; Garelli et al, 2012; Hackney & Cherbas, 2014; Halme et al, 2010).

However, slowing down peripheral growth poses a challenge to the regenerating cell population, which must continue to support high anabolic activity despite reduced canonical Insulin/PI3K/AKT signaling. The regenerating cell population circumvents this limitation by engaging a novel mechanism to sustain protein synthesis. Specifically, we find that Pdk1 is robustly upregulated in the proliferative domain of *egr-*expressing discs, and this upregulation is sufficient and necessary to drive ribosomal S6 phosphorylation and activate protein translation, even in the context of low systemic Insulin signaling. The capacity of Pdk1 to function independently of insulin positions Pdk1 as a central regulator of regenerative metabolism. Notably, our results mirror earlier observations of AKT-independent, yet Pdk1-dependent, growth during *Drosophila* development (Radimerski et al, 2002; Rintelen et al, 2001). During regeneration, we identify JAK/STAT as an upstream regulator of Pdk1, and consequently as a regulator for phosphorylation of ribosomal S6 protein and activation of protein synthesis. Our work thus uncovers a pathway whereby JAK/STAT signaling can mediate the known proliferation-promoting function in imaginal disc development, regeneration, and tumor growth (Herrera & Bach, 2019). Of note, the increase in Pdk1 protein levels likely arises from JAK/STAT regulated post-translational mechanisms affecting Pdk1 stability, as we did not observe an increase in *Pdk1* transcripts in *egr-*expressing discs (Floc’hlay et al, 2023). Such post-translational regulation of Pdk1 may also underlie the differential regulation of Pdk1 between regenerative and peripheral tissues. While we find that the proliferative domain actively upregulates Pdk1 to overcome insulin restrictions, we also find that the peripheral imaginal discs downregulate Pdk1. Based on our experiments demonstrating that Pdk1 levels are limiting for protein translation, the peripheral reduction in Pdk1 may contribute to the reduction in protein synthesis and growth of other imaginal discs. This dual regulation emphasizes the surprisingly central role of Pdk1 in integrating local demands for regeneration with systemic constraints on growth.

Furthermore, our work reveals that fat body catabolism, which mobilizes nutrients necessary for regeneration, is activated in *egr*-expressing larvae. In addition to remodeling of the lipid stores, proteins may also be catabolized, as suggested by an altered amino acid signature in the hemolymph. The upregulation of amino acid and carbohydrate transporters in the proliferative domain further suggests that regenerating cells actively import these mobilized nutrients, thereby fueling mTORC1 activation and subsequent protein synthesis. Earlier studies find elevated levels of ornithine, glutamate, and glutamine in rat wound fluids (Albina et al, 1993), similar to the ornithine, glutamate, and glutamine signature in hemolymph from egr-expressing larvae. Our observations suggest that glutamate metabolism may be important for regeneration, consistent with its described roles in tumor growth, inflammation, and defense against oxidative stress (Altman et al, 2016; Ling et al, 2023; Zhu & Thompson, 2019).

Together, our results support a model in which *egr*-induced inflammation induces systemic insulin restriction and insulin resistance, thereby limiting resource availability in peripheral tissues. At the same time, local activation of pathways such as JAK/STAT, Pdk1, or nutrient importers prioritizes tissue repair at the damage site. A potential limitation of our study lies in the high levels of *eiger-*expression used in this genetic model, which may not fully reflect physiological conditions. In fact, as we previously reported, *egr*-expressing discs can resemble chronic wound states, where prolonged inflammatory signaling impairs effective tissue regeneration (Jaiswal et al, 2023). However, in the widely studied *Ras^V12^,scrib-RNAi* tumor model, we observe that regions with high JAK/STAT activity also exhibit elevated Pdk1 expression and protein translation, suggesting that tumors which can induce cachexia also exploit this pathway to sustain growth despite low systemic insulin signaling. Given that Pdk1 can bypass the need for Insulin/AKT/FOXO signaling, it may not be surprising that Pdk1 upregulation is found in many cancers (Zheng et al, 2023). While oncogenic mutations may facilitate nutrient use in such a cachexic tumor environment (Lee et al, 2021; Newton et al, 2020; Santabarbara-Ruiz & Leopold, 2021; Song et al, 2019), our work specifically reveals how wild type tissues adapt within a physiological repair program to overcome systemic inhibition of growth.

In summary, our study uncovers a regulatory network wherein fat body catabolism, enhanced nutrient uptake, mTORC1 activation, and the instructive JAK/STAT-Pdk1 axis converge to support regenerative proliferation in cachexic environment. Undoubtedly, this is an ancient stress response and repair program designed to distribute nutrients from an energy store to the site of tissue damage. These findings not only advance our understanding of tissue repair mechanisms in *Drosophila* but may also offer insights into conserved pathways in mammals, as a domain characterized by ribosomal S6 protein phosphorylation was recently described in mouse and pig skin wounds, suggesting potential similarities in regenerative programs across species (Ring et al, 2023).

## Materials and Methods

### *Drosophila* maintenance

All experiments were performed on *Drosophila melanogaster*. Fly strains (see **Table S1**) were maintained on standard fly food (10L water, 74,5g agar, 243g dry yeast, 580g corn flour, 552ml molasses, 20.7g Nipagin, 35ml propionic acid) at 18°C – 22°C. Larvae from experimental crosses were allowed to feed on Bloomington formulation (175.7g Nutry-Fly,1100ml water 20g dry yeast, 1.45g Nipagin in 15ml Ethanol, 4.8ml Propionic acid) and raised at 18°C or 30°C to control GAL80ts-dependent induction of GAL4/UAS. Our experimental design did not consider differences between sexes unless for genetic crossing schemes.

### Drosophila genetics

To induce expression of UAS-constructs, such as *UAS-egr*, under the control of rn-GAL4 in the wing pouch, experiments were carried out as described in (Cosolo et al, 2019; La Fortezza et al, 2016; Smith-Bolton et al, 2009) with minor modifications. Briefly, larvae of genotype *rn-GAL4, tub-GAL80^ts^* and carrying the desired *UAS-*transgenes were staged with a 6h egg collection and raised at 18°C at a density of 50 larvae/vial. Overexpression of transgenes was induced by shifting the temperature to 30°C for 24h on the seventh day (D7) after egg deposition (AED) to relieve temperature-sensitive GAL80ts repression of GAL4. Larvae were dissected after 24 h of *egr-*expression. Control genotypes were generated by crossing rn-GAL4, tub-GAL80ts into a wild type background. The DUAL Control genetic ablation system (DCS) was employed, wherein a single heat shock simultaneously activates genetic cell ablation by egr-expression within the *salm* domain and GAL4 expression in the *dve-GAL4* domain (*hsFLP; hs-p65::zip, lexAOp-egrNI/CyO; salm-zip::LexA-DBD, DVE>>GAL4)* (Harris et al, 2020). Specifically, larvae were raised at 25°C and subjected to a 1-hour heat shock at 37°C on Day 4.5, followed by dissection 24 hours post-heat shock. The Pdk1-GFP line (BDSC: 59836) was characterized using the following approaches: a complex expression pattern can be observed in wild type tissues in immunofluorescence, where GFP is detected in the cytoplasm but also at membranes (see for example, Fig 6C). Tissues form larvae produce a band of the expected size on Western blot. Three independent RNAi lines (BDSC: 27725, BDSC: 34936, BDSC: 36071) targeting the 3’ region of the Pdk1 transcript downstream of the GFP-cassette insertion site can robustly knock-down GFP expression (see for example Fig 5E). We conclude that a full-length, membrane-recruitable and genetically tractable protein is produced from the endogenous locus. Thus. the annotation of the insertion site in FlyBase is incorrectly oriented.

### Immunohistochemistry of wing imaginal discs

Wing discs from third instar larvae were dissected and fixed for 15 minutes at room temperature in 4% paraformaldehyde in PBS. Washes were performed in PBS containing 0.1% TritonX-100 (PBT). The discs were then incubated with primary antibodies (listed in **Table S2**) in 0.1% PBT, gently mixed overnight at 4°C. During incubation with cross-absorbed secondary antibodies coupled to Alexa Fluorophores at room temperature for 2 hours, tissues were counterstained with DAPI (0.25 ng/µL, Sigma, D9542). Tissues were mounted using SlowFade Gold Antifade (Invitrogen, S36936). To ensure comparability in staining between different genotypes, experimental and control discs were processed in the same vial and mounted on the same slides whenever possible. Images were acquired using the Leica TCS SP8 Microscope, using the same confocal settings for linked samples and processed using tools in Fiji.

### Protein synthesis assays using OPP-Click-iT staining

OPP Assays were performed using Click-iT® Plus OPP Protein Synthesis Assay Kits (Invitrogen Molecular Probe) according to manufacturer’s instructions. Briefly, larvae were dissected and inverted cuticles were incubated with a 1:1000 dilution of Component A in Schneider’s medium for 15 minutes on a nutator. Larval cuticles were fixed with 4% paraformaldehyde for 15 minutes, rinsed twice in 0.1% PBT, and permeabilized with 0.5% PBT for 15 minutes. The cuticles were then stained with the Click-iT® cocktail for 30 minutes at room temperature, protected from light. Further immunohistochemistry analysis and sample mounting was performed as described above.

### EdU labelling

EdU incorporation was performed using the Click-iT Plus EdU Alexa Fluor 647 Imaging Kit. Briefly, larval cuticles were inverted in Schneider’s medium and incubated with EdU (10µM final concentration) at RT for 15 minutes. Cuticles were then fixed in 4% PFA/PBS for 15 minutes, washed for 30 minutes in PBT 0.5%. EdU-Click-iT labeling was performed according to manufacturer’s guidelines. Further immunohistochemistry analysis and sample mounting was performed as described above.

### SA-β-Gal staining

Cell senescence detection kit from Invitrogen (C10850) was used to analyze senescence-associated β-galactosidase activity. Briefly, larval cuticles were inverted in PBS, fixed with 4% PFA, washed with 1% BSA (in PBS) and then incubated in working solution for 2h at 37°C, according to manufacturer’s instructions. Washing steps were performed in PBS and PBS containing 0.1% TritonX-100 (PBT). Further immunohistochemistry analysis and sample mounting was performed as described above.

### Fat body Nile Red staining

Early third instar larvae were collected in PBS and dissected, leaving the gut intact to prevent fat body loss. Inverted cuticles were transferred to an Eppendorf tube and fixed with 4% paraformaldehyde/PBS for 15 minutes. Samples were washed in 0.1% PBT. Cuticles were incubated with Nile Red (2 μg/mL in PBS) for 1 hour, protected from light. Following incubation, samples were washed in PBS, the fat body was dissected and mounted as described above.

### Hemolymph sample preparation

Fifteen larvae were collected and thoroughly washed with Milli-Q water to remove any fly food particles. Care was taken to ensure no food particles remained on the larvae surfaces, and they were dried using Kim Tech paper wipes. Each larva was punctured in the center using forceps and transferred to a 0.5 mL microcentrifuge tube with three 1 mm holes at the bottom. The 0.5 mL microcentrifuge tube was then placed into a pre-cooled 1.5 mL microcentrifuge tube. One larva at a time was processed in this assembly and centrifuged in a microfuge for 10 seconds. After each centrifugation, the larval carcass was removed from the 0.5 mL microcentrifuge tube to prevent blockage of the holes. Hemolymph isolated from the 15 larvae was collected in the bottom 1.5 mL tube. From the total collected hemolymph, 8 µL was transferred to a fresh 1.5 mL tube, and 10 µL of ultrapure Milli-Q water, 30 µL of methanol, an internal standard, and 50 µL of MTBE were added. The solution was mixed thoroughly and centrifuged at 1000g for 10 minutes at 4°C. After centrifugation, the organic and polar phases were collected separately in different tubes for metabolite measurement.

### Hemolymph metabolomic analyses

Non-targeted analysis of polar metabolites by LC-MS was carried out as described previously (Edwards-Hicks et al, 2020) using an Agilent 1290 Infinity II UHPLC in line with a Bruker Impact II QTOF-MS operating in negative and positive ion mode. Scan range was from 20 to 1050 Da and mass calibration was performed at the beginning of each run. LC separation was on a Waters Atlantis Premier BEH ZHILIC column (100 x 2.1 mm, 1.7 µm particles), buffer A was 20 mM ammonium carbonate and 5 µM medronic acid in milliQ H2O and buffer B was 90:10 acetonitrile:buffer A and the solvent gradient was from 95% to 55% buffer B over 14 minutes. Flow rate was 180 µL/min, column temperature was 35 °C, autosampler temperature was 5 °C and injection volume was 3 µL. Data processing including feature detection, feature deconvolution, and annotation of features was performed using MetaboScape (version 2023b). Quantile normalization was performed to minimize sample size effects and further statistical processing were performed with R. Metabolites with missing values were eliminated from the dataset. Both, the Shapiro-Wilk test and the Q-Q plot, showed that the data was not normally distributed. Therefore, a two-sided, unpaired Wilcoxon rank-sum test was performed on the quantile normalized data. Only metabolites with at least <0.75 and >1.5-fold change and a p-value < 0.19 and were selected for further analysis and were displayed in a heatmap with row-wise normalization (Fig 3L).

### Thin Layer Chromatography (TLC)

Fat bodies were dissected in cold PBS from third instar (L3) control (rn-GAL4, tubGAL80ts) and egr-expressing (rn-GAL4, UAS-egr, tubGAL80ts)larvae, and from wild type larvae starved for 24 h. Three fat bodies were pooled per sample and immediately transferred into 100μL chloroform: methanol (3:1) solution, followed by storage on ice. Samples were mechanically homogenized and centrifuged at 15,000 × g for 5 minutes at 4°C. For semi-quantitative analysis of triglycerides, a standard curve was prepared using lard as a reference material, dissolved in chloroform: methanol (3:1) at the following concentrations: STD1 (75 μg/μL); STD2 (60 μg/μL); STD3 (50 μg/μL); STD4 (37.5 μg/μL); STD5 (25 μg/μL); STD6 (15 μg/μL); STD7 (10 μg/μL). A blank control containing only the solvent mixture (chloroform:methanol (3:1)) was included. Samples and standards were loaded onto a glass silica gel TLC plate (Millipore). The mobile phase consisted of a hexane:diethyl ether (4:1) solvent system. Chromatography was performed and after air-drying, the TLC plate was treated with ceric ammonium heptamolybdate (CAM) staining solution and developed at 80°C, with band development monitored at 5-minute intervals. Images were captured using a gel documentation system (gelONE); image analysis and quantification were performed using ImageJ. Automatic image thresholding was applied using the Otsu method, and regions of interest (ROIs) were defined for each band. Integrated intensity analysis was performed, and triglyceride density was quantified based on the standard curve.

### Real-Time PCR

Third instar (L3) control (rn-GAL4, tubGAL80ts), egr expressing (rn-GAL4, UAS-egr, tubGAL80ts), and wild type larvae starved for 24 h larvae were carefully collected from the food and pooled in groups of three male larvae per sample. Samples were homogenized in 100 µL of TRIzol, and RNA was extracted using chloroform (20 µL), followed by precipitation with 50 µL of isopropanol. The purified RNA was suspended in nuclease-free water, subjected to DNase treatment and reverse transcription was performed using RevertAid Reverse Transcriptase (Thermo Fisher Scientific) according to manufacturer’s guidelines. qPCR was performed using the Blue S’Green qPCR Kit Separate ROX (Biozym) on a LightCycler 480 (Roche). Standard dilution series (1:5) were generated for each primer pair to determine primer efficiency based on the regression curve. The following primers were used (for, rev 5’-3’): Rpl1 (TCCACCTTGAAGAAGGGCTA, TTGCGGATCTCCTCAGACTT), Ilp2 (ATCCCGTGATTCCACCACAAG, GCGGTTCCGATATCGAGTTA), Ilp5 (GCCTTGATGGACATGCTGA, AGCTATCCAAATCCGCCA). Gene expression levels were normalized to the reference gene Rpl1, and relative expression was calculated using the Pfaffl method. Statistical outliers were identified using the ROUT test (Q = 1%) and were removed to maintain data consistency. Statistical analysis was performed using the Kruskal-Wallis test.

### Western Blot Analysis

Fat bodies were dissected in cold PBS from third-instar (L3) control (rn-GAL4, tubGAL80ts) and egr-expressing (rn-GAL4, UAS-egr, tubGAL80ts) larvae, and from wild type larvae starved for 24 h. Five fat bodies were lysed on ice (300mM NaCl, 50mM Tris-HCl ph 7,5, 1% Triton X-100, 0.1mM EDTA, 0.1% SDS, 5% Glycerol, Roche Complete protease inhibitor cocktail and Complete phosphate inhibitor cocktail). Tissues were further homogenized and centrifuged at 6000g for 3 minutes at 4°C. Supernatants were incubated in Laemmli buffer at 85°C for 5 minutes. Samples were loaded onto Mini-PROTEAN TGX gels (4–15%) corresponding to one fat body per lane. After electrophoresis, proteins were transferred onto a nitrocellulose membrane, blocked in 1× Tris-buffered saline containing bovine serum albumin, incubated overnight with primary antibodies at 4°C and secondary antibody for 1 h at RT. Proteins were visualized using SuperSignal West Femto Maximum Sensitivity Substrate and a Bio-Rad ChemiDoc-MP imaging system.

### Image quantification and statistical analysis-General comments

For all quantifications, control and experimental samples were processed together and imaged in parallel, using the same confocal settings. Images were processed, analyzed and quantified using tools in Fiji (ImageJ 2.14) (Schindelin et al, 2012). Extreme care was taken to apply consistent methods to control and experimental samples (i.e., number of projected sections, thresholding methods, processing) for image analysis and quantifications. See **File S1** for macros used in FIJI during image segmentation and quantifications. Further details on segmentation and quantifications are provided below. Figure panels were assembled using Affinity Designer 2.4.0. Statistical analyses were performed in Graphpad Prism. Illustrations were done in Biorender.

To quantify fluorescence intensities inside the proliferative domain (PD^egr^), a mask of the high JNK-signaling domain region was generated, followed by creating a 20μm band on the outside to mark the proliferative domain. Fluorescence intensities in proliferative domains was always compared to fluorescence intensities within the pouch domain of control discs, unless otherwise noted.

### Quantification of cycling cells using EdU incorporation

#### (a) EdU quantification in notum

An xy-section with the maximum number of notum epithelial cells was selected, excluding any myoblast cells. A mask was created from the DAPI staining to represent nuclei and saved as ROI. A binary operation (AND) was used to compute a notum DAPI mask with the aid of a manually selected notum region (FileS1, Macro-1L(a)). This mask was then used to measure EdU intensity.

#### (b) EdU quantification within 10μm bands

In *egr-*expressing discs, TRE-RFP expression was used to generate a mask of the high JNK-signaling domain (FileS1, Macro-1D(a)). The boundary of this mask defined the regions inside and outside the high JNK region. Five 10μm bands were created outside the high JNK-signaling domain, and three 10μm bands inside of the JNK-signaling domain. Also applying a DAPI mask to each band (FileS1, Macro-1D(b)), the mean EdU intensity in each of the 10μm bands was measured. Consistent with previous studies (Cosolo et al, 2019; Jaiswal et al, 2023), the highest EdU intensities were observed in 3 bands outside the JNK-signaling domain, which was denoted as proliferative domain.

#### (c) EdU quantification in eye disc

For the eye disc, an EdU mask (FileS1, Macro-2T(a)) was generated using the EdU staining, which was then applied to measure EdU intensity in the entire visible eye disc.

### Quantification of Upd3-LacZ levels

A maximum projection of TRE-RFP expression was used to generate a mask of the high JNK-signaling domain in *egr-*expressing disc (FileS1, Macro-S3H(a)). Upd3-LacZ intensity was measured within the TRE-RFP mask of the high JNK-signaling domain in *egr-*expressing disc, while for the control disc, Upd3-LacZ intensity was measured in the pouch.

### Quantification of ImpL2-GFP levels

A pouch mask was generated using the maximum projection of an anti-Nubbin staining (FileS1, Macro-3H(a)) for both control and *egr-*expressing discs. ImpL2-GFP intensities were measured within the Nubbin mask on the sum projection of the ImpL2-GFP signals within the stack (seven slices in each disc).

### Quantification of OPP levels

#### (a) in different regions of wing imaginal disc

In the control disc, mean OPP intensity was measured separately in the pouch, hinge, and notum regions. In *egr-*expressing discs, mean OPP intensity was measured in the high JNK signaling region (TRE-RFP mask), the proliferative region (20µm band around the TRE-RFP mask), and the notum region. In control discs, the pouch, hinge, and notum regions were selected using wing fold landmarks. In *egr-*expressing discs, a max projection of TRE-RFP intensities was used to create a mask of the high JNK signaling domain (FileS1, Macro-1G(a)), which was applied to measure OPP intensity in the high JNK region. A 20μm band outside the high JNK region was generated to mark the proliferative region, and OPP intensity in this domain was measured. For the notum, a manual selection along the first notum fold and outlining the edges of the notum was carried out.

#### (b) in the eye disc

For the eye-antennal disc, the DAPI staining was used to generate a mask of the tissue outline (FileS1, Macro-2Q(a)) and combined with a manual selection of the eye disc to generate a region of interest (ROI) within which we measured OPP intensity.

### Quantification of nuclear dFOXO levels

In control discs, a slice with the maximum number of pouch cells was selected and a DAPI mask (FileS1, Macro-4G(a)) was generated and combined with manual pouch selection to create a pouch nuclear mask. This mask was used to measure mean dFOXO intensity in the pouch nucleus. In *egr-*expressing discs, a single slice from the Z-stack with the maximum number of cells in the proliferative domain was selected. A max projection of TRE-RFP intensities was used to create a mask of the high JNK signaling domain, and a 20μm band was created to locate the proliferative region. A DAPI mask (FileS1, Macro-4G(b)) was generated from the selected slice and combined with the 20μm band to create a new mask for the proliferative cell nuclei, which was used to measure dFOXO intensity in the nucleus. For the analysis of nuclear dFOXO in the notum, a Z-slice containing the maximum number of notum cells was selected. A DAPI mask (File S1, Macro-2F(a)) was generated and combined with manual selection of notum, guided by wing fold, to create a notum nuclear mask. Please note that discs and fat body were treated with Leptomycin B (413nM) for 30 min before dissection to block nuclear export.

### Quantification of Pdk1 GFP, p-S6 and OPP levels in the en-GAL4 domain

The slice with the maximum number of pouch cells was selected, and masks of the anterior and posterior pouch compartments were generated by segmenting the enGAL4, UAS-RFP expression domain. Levels of p-S6(File S1, Macro 6F(a)) and Pdk1-GFP intensities were measured within these masks and the P/A ratio (en-GAL4 region/non-en-GAL4 region) was calculated. Mean OPP intensity in the control and experimental pouch en-GAL4 domain was measured and reported.

### Quantification of 10xStat92E-GFP levels in the fat body

A single slice from the Z-stack with the maximum number of nuclei was selected. A 200×200 pixel square region was defined and used as a standard area for measurement of GFP intensities.

### Quantification of lipid droplet size and shape

A single slice from the Z-stack through the most anterior fat body containing the maximum number of nuclei was selected. A published STAR protocol was applied to measure lipid droplet area and the ‘roundness’ of lipid droplets (defined as the relationship between area and length of major droplet axis) in both control and experimental samples (Dark et al, 2022).

## Data availability

All data, workflows and FIJI based algorhithms necessary to interpret the data are included within the manuscript. Original microscopy image files and intermediate segmentation results will be made available upon request. Requests should be addressed to and will be fulfilled by AKC.

## Inclusion & ethics statement

All authors fulfill the criteria to be included as co-authors. All authors agreed to their roles and responsibilities in the study design.

## Author Contributions

Conceptualization AVM, AKC; Data curation AVM, LNe, LNo, LH, AKC; Formal analysis AVM, LNe, LH, JB, AKC; Funding acquisition KK, AKC; Investigation AVM, LNe, LNo, AS, AKC; Methodology AVM, LNe, LNo, AS, IG, JB, AKC; Project administration IG, AKC; Supervision KK, AKC; Validation AVM, LNe, LNo, AS, KK, AKC; Visualization AVM, LNe, LH; Writing – original draft AVM, AKC; Writing – review & editing AVM, KK, AKC

## Acknowledgements

We thank the staff of the Life Imaging Center (LIC) in the Hilde Mangold House (HMH) of the Albert-Ludwigs-University of Freiburg for help with their confocal microscopy resources, and the excellent support in image recording. We specifically thank the DFG for supporting our imaging work through project number 414136422. We thank the Lighthouse Core Facility and its staff, as well as Elisabeth Savkova, Janhvi Jaiswal and Carlo Crucianelli for help with our experimental work. We thank Robin Harris, Susumi Hirobayashi, Hugo Stocker, Aurelio Teleman and Hong Xu, Iswar Hariharan, Dirk Bohmann for sharing reagents and the Bloomington *Drosophila* Stock Center (BDSC), the Vienna *Drosophila* Stock Collection (VDRC), the University of Zurich ORFeome Project (FlyORF), the Developmental Studies Hybridoma Bank (DSHB) and the Monoclonal Antibody Core Facility at the Helmholtz Zentrum Munich for providing fly stocks and antibodies. We thank the IMPRS-EBM and SGBM graduate schools for supporting our doctoral researchers.

## SUPPLEMENTARY FIGURE LEGENDS

**Figure S1.**
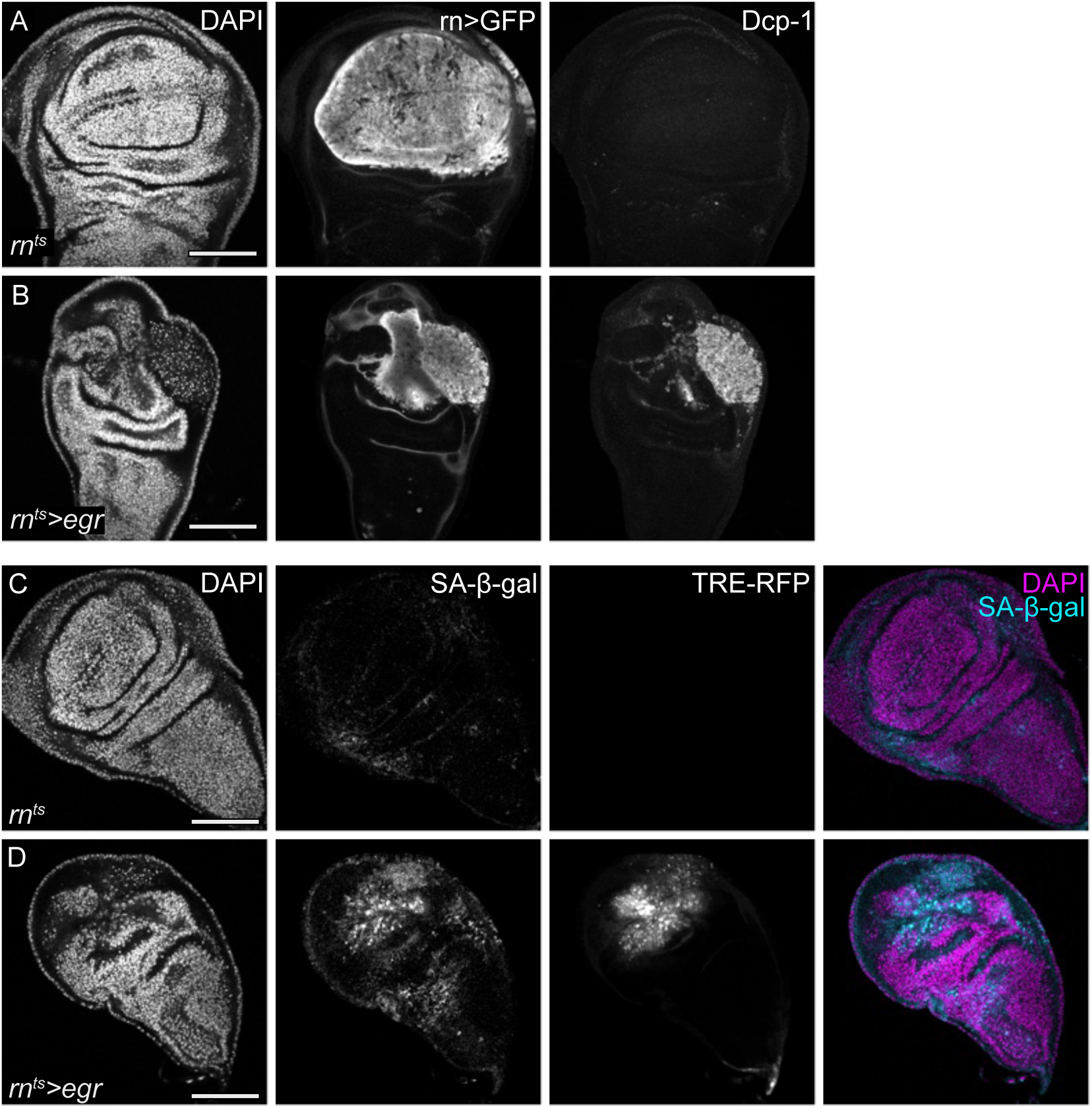
Cell cycle adaptations in response to inflammatory tissue damage. **A, B.** Control wing disc (A) and a wing discs (B) after 24 hours of *egr-*expression within the wing pouch domain under the control of the *rn*-GAL4 driver (*rn^ts^* and *rn^ts^>egr*, respectively). A co-expressed UAS-GFP construct visualizes the *rn*-GAL4 expression domain. Staining for cleaved Dcp-1 visualizes apoptosis. Discs were stained with DAPI to visualize nuclei. Please note the GFP-expression domain in (B) which does not stain for cDcp-1 and represents surviving cells with high levels of JNK activity. Levels of Dcp-1 were previously quantified (La Fortezza, Schenk et al. 2016, Cosolo, Jaiswal et al. 2019, Jaiswal, Egert et al. 2023). **C, D.** Analysis of senescence-associated β-galactosidase (SA-β-gal) activity (cyan or grey) in control (C) and *egr-*expressing wing disc (D). TRE-RFP visualizes JNK-pathway activity. Discs were stained with DAPI to visualize nuclei (magenta or grey). SA-β-gal activity was already previously reported and quantified (Floc’hlay, Balaji et al. 2023, Jaiswal, Egert et al. 2023). Scale bars: 100 μm.

**Figure S2.**
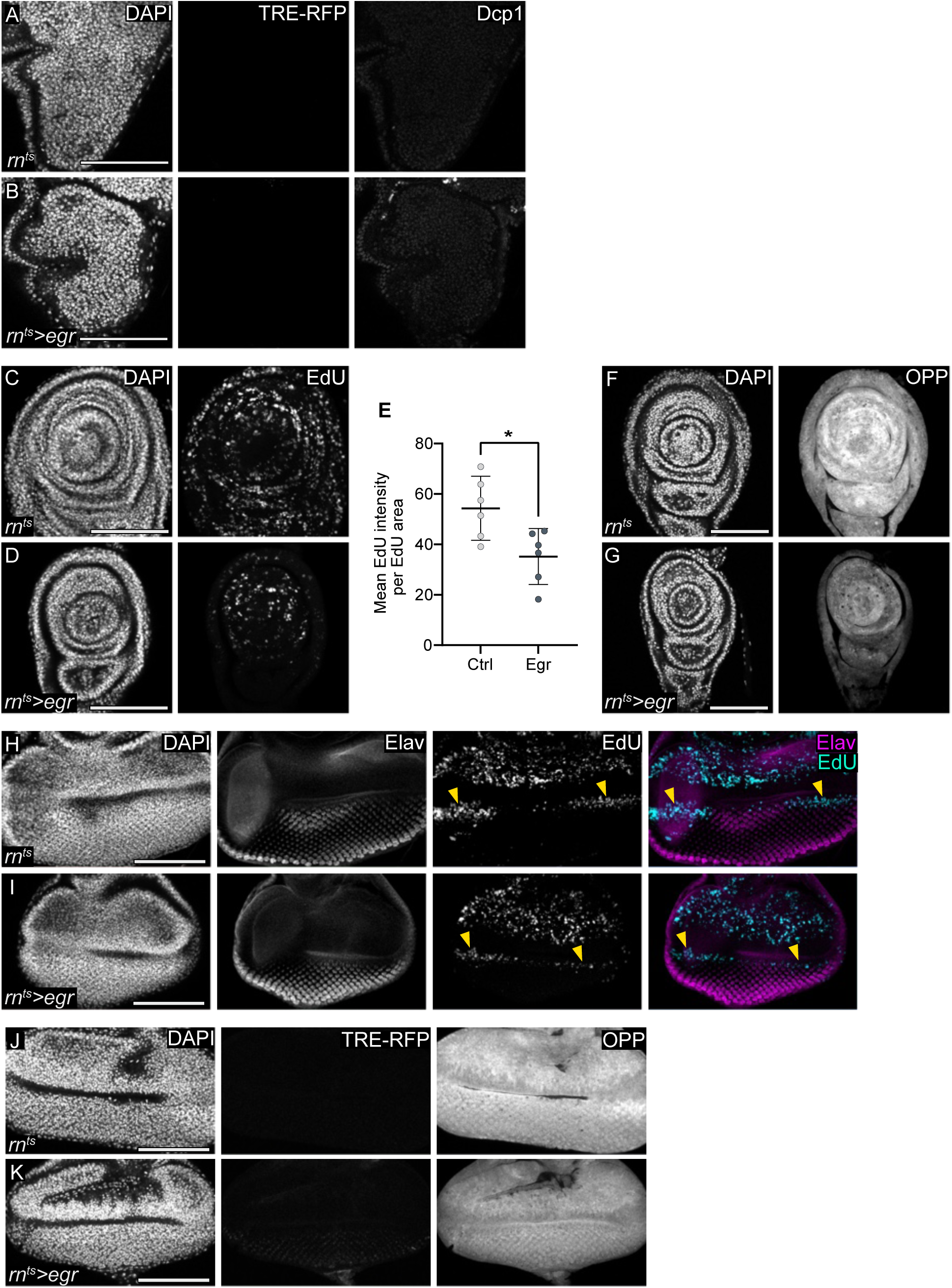
Metabolic adaptation in peripheral tissues. **A, B.** Nota of control (A) and wing disc expressing *egr* in the pouch (B). TRE-RFP visualizes JNK-pathway activity and staining for cleaved Dcp-1 visualizes apoptosis. Discs were stained with DAPI to visualize nuclei. **C, D.** EdU incorporation to visualize DNA replication in leg imaginal discs, dissected from larvae containing either control wing disc (C) or *egr-*expressing wing discs (D). Discs were stained with DAPI to visualize nuclei. **E.** Mean EdU intensity per EdU-positive area quantified in leg imaginal discs, dissected from larvae containing either control wing disc (C) or *egr-*expressing wing discs (D), serving as a proxy for DNA replication speed. Mean and 95% CI are shown. Statistical significance was tested using an two-tailed Unpaired t-test with a p-value = 0.0154 (control: n=6, *egr-*expressing disc: n=6). **F, G.** Protein synthesis visualized by O-proparyl-puromycin (OPP) incorporation into newly synthesized proteins in leg imaginal discs, dissected from larvae containing either control wing disc (E) or *egr-*expressing wing discs (G). Discs were stained with DAPI to visualize nuclei. **H, I.** EdU incorporation (cyan or grey) to visualizes DNA replication in eye imaginal discs, dissected from larvae with control (H) or *egr-*expressing (I) wing imaginal discs. Eye disc differentiating photoreceptors are marked with Elav staining (magenta or grey). Sum projection of multiple image slices to visualize the second mitotic wave (SMW) and the yellow arrowheads point to the expected location of the SMW. **J, K.** Protein synthesis visualized by OPP incorporation in the eye imaginal disc (same as Fig 2**.O and P**), dissected from larvae with control (J,) or *egr-*expressing (K) wing imaginal discs using *rn*-GAL4 driver. TRE-RFP visualizes JNK-pathway activity and discs were stained with DAPI to visualize nuclei. Scale bar: 100 μm.

**Figure S3.1.**
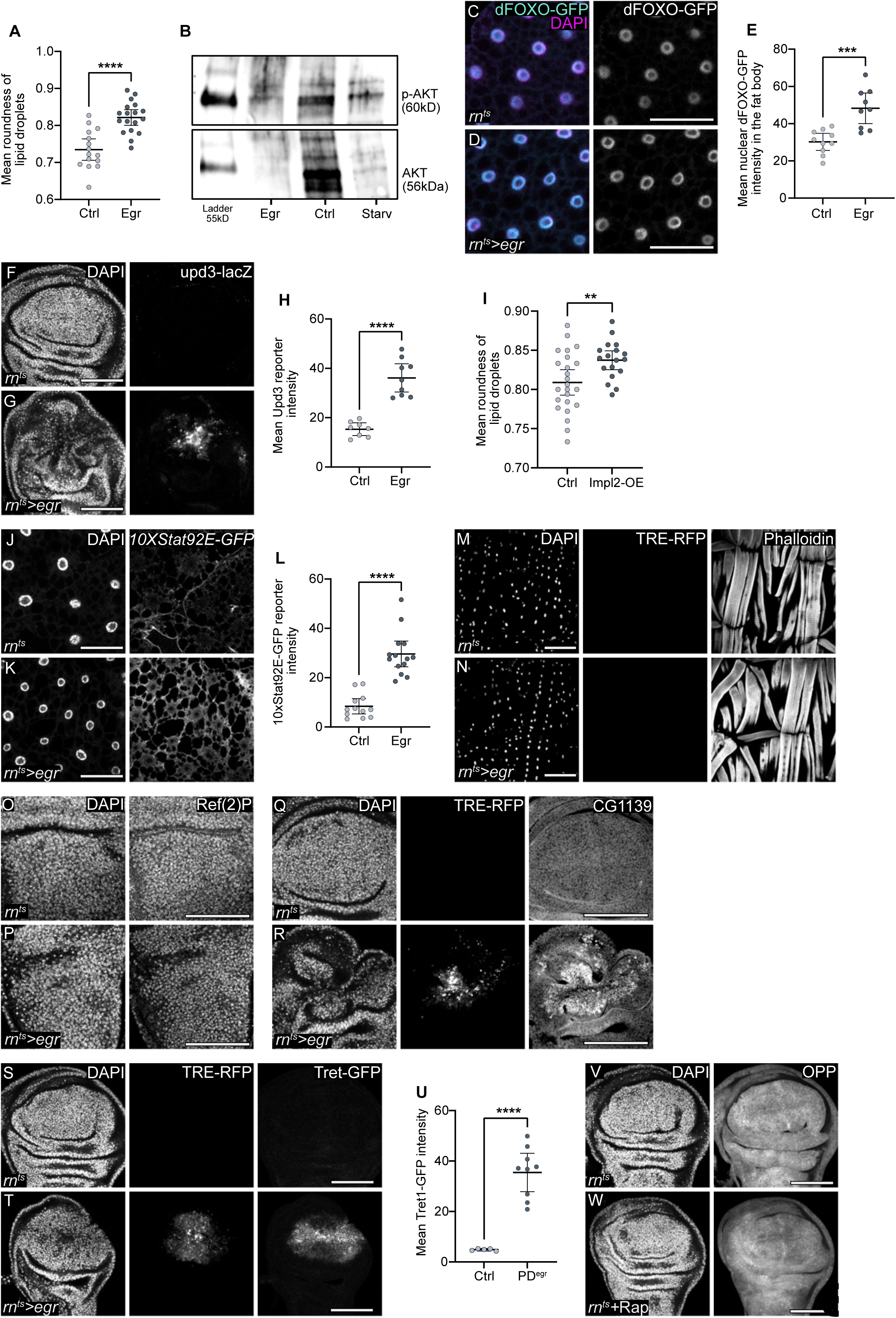
Elevated levels of nutrient transporters and mTORC1 signaling support regenerative proliferation. **A.** Quantification of roundness of lipid droplets in the fat body dissected from larvae with control (Fig.3A) or *egr-*expressing (Fig.3B) wing imaginal discs. Mean and 95% CI are shown. Roundness is measured as the ratio of the area to the square of the length of the major axis. A perfect circle has a roundness value of 1, while values approaching zero indicate a more elongated shape. Statistical significance of was tested using an two-tailed Unpaired t-test. Mean roundness, p-value < 0.0001. (control: n=15, *egr-*expression in discs: n=18). **B.** Protein expression levels of p-AKT (Ser473) and AKT by western blot in the fat body dissected from larvae with control or *egr-*expressing wing imaginal discs or 24 h starved larvae. **C, D.** Expression of dFOXO-GFP (cyan or grey) in fat body, dissected from larvae with control (I) or *egr-*expressing (J) wing imaginal discs. Fat bodies were stained with DAPI to visualize nuclei (magenta). **E.** Mean nuclear dFOXO-GFP intensity in fat body, dissected from larvae with control or *egr-*expressing wing imaginal discs. Statistical significance was tested using an two-tailed unpaired t-test, p-value = 0.0003 (control: n=10, *egr-*expressing disc: n=9). **F, G**. Expression of Upd3-lacZ in control (C) and *egr-*expressing discs (D). Discs were stained with DAPI to visualize nuclei. **H.** Upd3-lacZ intensity quantified in the pouch of control wing discs and in the JNK-signaling domain of *egr-*expressing discs. Mean and 95% CI are shown and statistical significance was tested using two-tailed Welch’s t-test with p-value < 0.0001 (control: n=8, *egr-*expressing disc: n=9). **I.** Quantification of roundness of lipid droplets in the fat body dissected from larvae with either control discs or wing imaginal discs expressing ImpL2 for 24 h under the control of *rn*-GAL4. Roundness is measured as the ratio of the area to the square of the major axis. A perfect circle has a roundness value of 1, while values approaching zero indicate more elongated shape. Statistical significance was tested using the two-tailed Unpaired t-test, p-value=0.0088 (control: n=24, *egr-*expression in disc: n=18). **J, K.** Activity of the JAK/STAT reporter *STAT92E-GFP* in the fat body dissected from larvae with control (J) or *egr-*expressing (K) wing imaginal discs. DAPI visualizes nuclei. **L.** Quantification of *STAT92E-GFP* reporter intensity in the fat body dissected from larvae with control or *egr-*expressing wing imaginal discs. Mean and 95% CI are shown and statistical significance was tested using the two-tailed Mann-Whitney test, p-value < 0.0001 (control: n=12, *egr-*expression in disc: n=14). **M, N.** *Drosophila* larval body wall muscles dissected from larvae containing either control (M) or *egr-*expressing (N) wing discs. Actin was visualized using Phalloidin and TRE-RFP visualizes JNK-pathway activation. Muscles were stained with DAPI to visualize nuclei. Please note that muscles do not activate JNK-signaling in response to inflammatory cues. **O, P.** Nota of control (O) and *egr-*expressing wing disc (P) stained for the autophagy marker Ref(2)P. Discs were stained with DAPI to visualize nuclei. **Q, R.** Staining for the putative alanine, glycine and proline amino acid transporter Arcus (CG1139) in control (Q) and *egr-*expressing discs (R). TRE-RFP visualizes JNK-pathway activation. Discs were stained with DAPI to visualize nuclei. **S, T.** Expression of Tret1.1-GFP in control (S) and *egr-*expressing discs (T). TRE-RFP visualizes JNK-pathway activation. Discs were stained with DAPI to visualize nuclei. **U.** Mean Tret1.1-GFP intensity quantified in the pouch of control discs and the proliferative domain of *egr-*expressing discs. Mean and 95% CI are shown. Statistical significance was tested using the two-tailed Welch’s t-test, p-value < 0.0001 (control: n=5, *egr-*expressing discs: n=9). **V, W.** Protein synthesis visualized by OPP incorporation in control (V) and larvae fed on food with rapamycin (200 μM) (W) for 24 h during the temperature shift. Scale bar: 100 μm. Fluorescence intensities are reported as arbitrary units.

**Figure S3.2.**
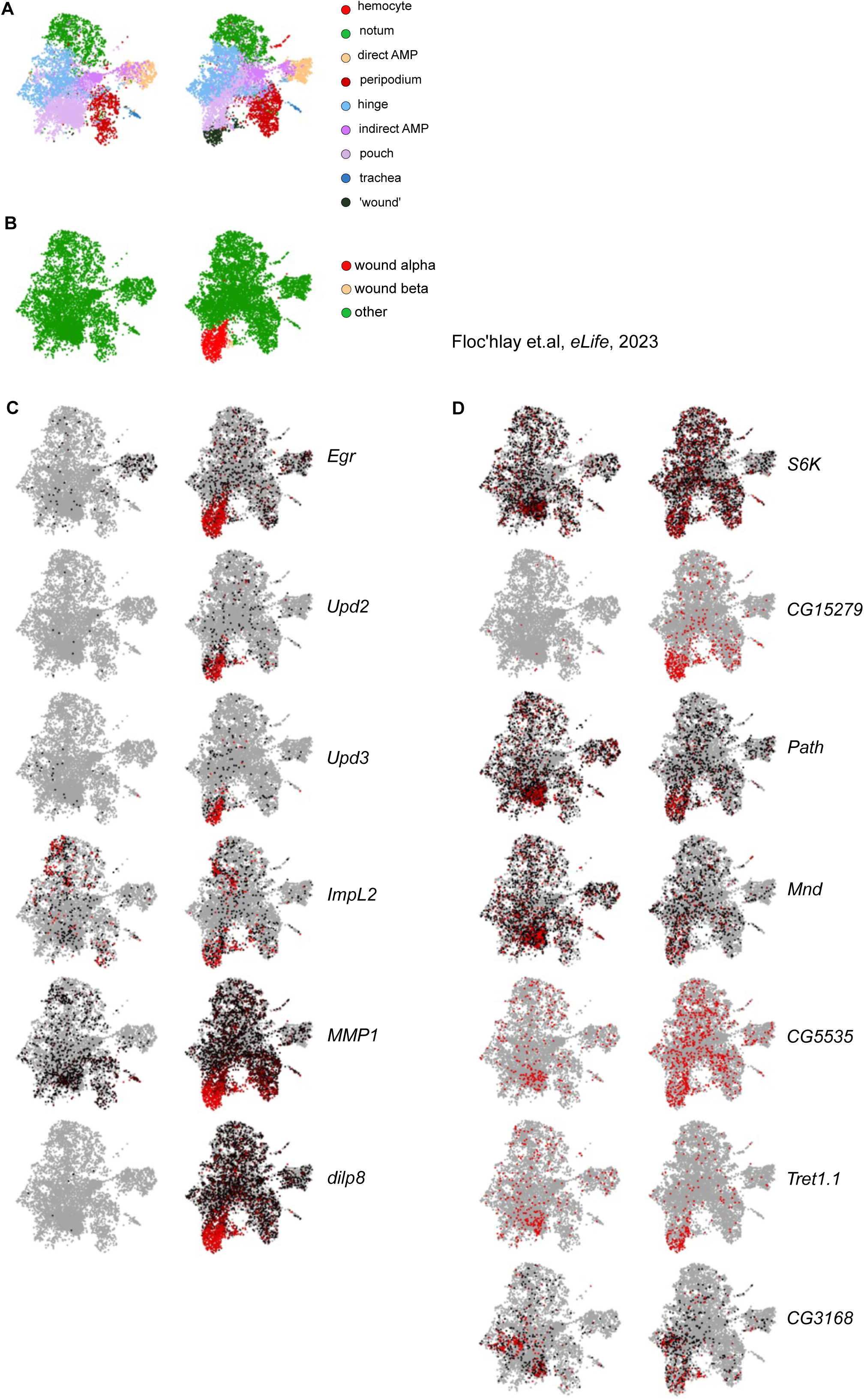
Elevated levels of nutrient transporters and mTORC1 signaling support regenerative proliferation. **A.B** UMAP plots generated using the Scope Wing Atlas created from control and *egr-*expressing discs analysed by single-cell RNA-Seq technology (Floc’hlay, Balaji et al. 2022). Different cell populations are color-coded. The analysis in (Floc’hlay, Balaji et al. 2022) indicating the emergence of a ‘wound’ cluster in *egr-*expressing discs, subdivided into two clusters with signatures characteristic of senescent cells (beta) and with characteristic of a cell and tissue damage response signature (alpha). **C.** ScRNA-Seq UMAP plots showing the elevated transcript levels of secreted paracrine ligands with known interorgan signaling roles. Grey indicates low expression, and red indicates high expression of transcripts. **D.** scRNA-Seq UMAP plot displaying significantly elevated transcript levels detected for different nutrient transporters and S6K (Floc’hlay, Balaji et al. 2023). Grey indicates low expression, and red indicates high expression of transcripts.

**Figure S4.**
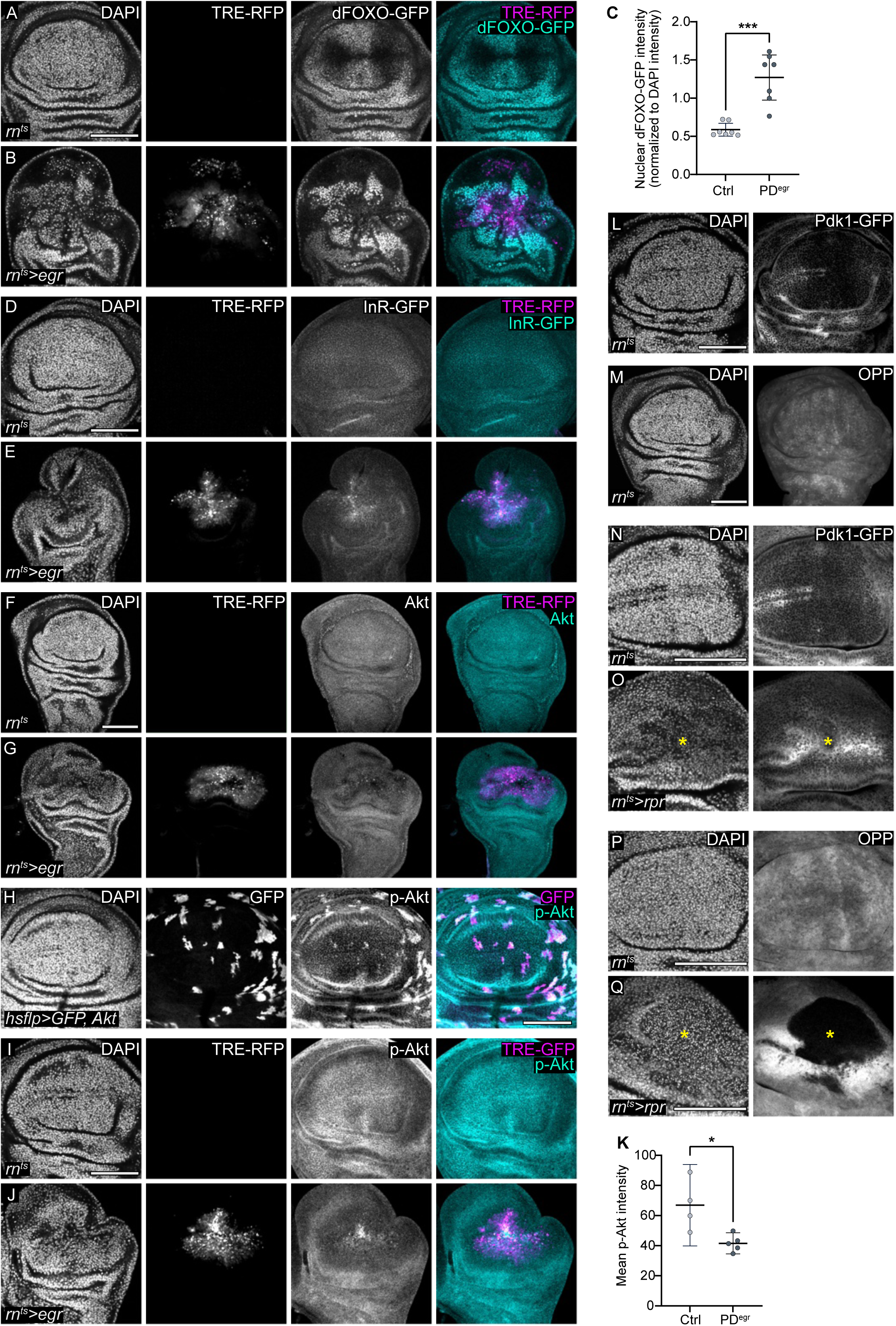
Insulin/PI3K/Akt signaling is low in the proliferative domain. **A,B.** Expression of dFOXO-GFP (cyan or grey) in control (A) and *egr-*expressing discs (B). TRE-RFP visualizes JNK-pathway activity (magenta or grey) and DAPI visualizes nuclei (grey). dFOXO-GFP expression is visualized using BDSC 59766 fly line. **C.** Mean nuclear dFOXO-GFP intensity quantified in the pouch of control discs or the proliferative domain of *egr-*expressing discs (PD^egr^). Mean and 95% CI are shown and statistical significance was tested using the two-tailed Mann-Whitney test, p-value = 0.0006 (control: n=7, *egr-*expressing disc: n=7). **D-G.** Control (D, F) and *egr-*expressing wing discs (E, G), expressing an Insulin receptor tagged with GFP (InR-GFP) (D, E) or stained for Akt (F, G) (cyan or grey). TRE-RFP visualizes JNK-pathway activity (magenta or grey). Discs were stained with DAPI to visualize nuclei. **H.** Anti-phospho-Akt (S505) staining (cyan or grey) to test antibody specificity in UAS-Akt-overexpressing clones marked by co-expression of UAS-GFP (magenta or grey) in the wing imaginal disc. Discs were stained with DAPI to visualize nuclei. **I, J.** Control (I) and *egr-*expressing wing discs (J), stained for phospho-Akt (S505) (cyan or grey). TRE-RFP visualizes JNK-pathway activity (magenta or grey). Discs were stained with DAPI to visualize nuclei. **K.** Quantification of mean p-AKT intensity in the control pouch and proliferative domain of *egr-*expressing wing discs. Mean and 95% CI are shown and statistical significance was tested using a two-tailed Unpaired t-test, p-value =0.0156. (Control; n=4, *egr-*expressing discs; n=5). **L.** Expression of Pdk1-GFP in the control wing imaginal disc corresponding to pro-apoptotic *hid-*expressing discs shown in **Fig4.K**. DAPI visualizes nuclei. **M.** Protein synthesis visualized by OPP incorporation in the control wing imaginal disc corresponding to *hid-*expressing discs shown in **Fig4.L**. DAPI visualizes nuclei. **N, O.** Expression of Pdk1-GFP in the control and pro-apoptotic gene Reaper (*rpr*) expressing discs. The yellow asterisk marks the damage or *rpr*-expressing region. DAPI visualizes nuclei. **P, Q.** Protein synthesis visualized by OPP incorporation in the control and pro-apoptotic gene Reaper (*rpr*) expressing discs. The yellow asterisk marks the damage or *rpr*-expressing region. DAPI visualizes nuclei. Scale bar: 100 μm.

**Figure S5.**
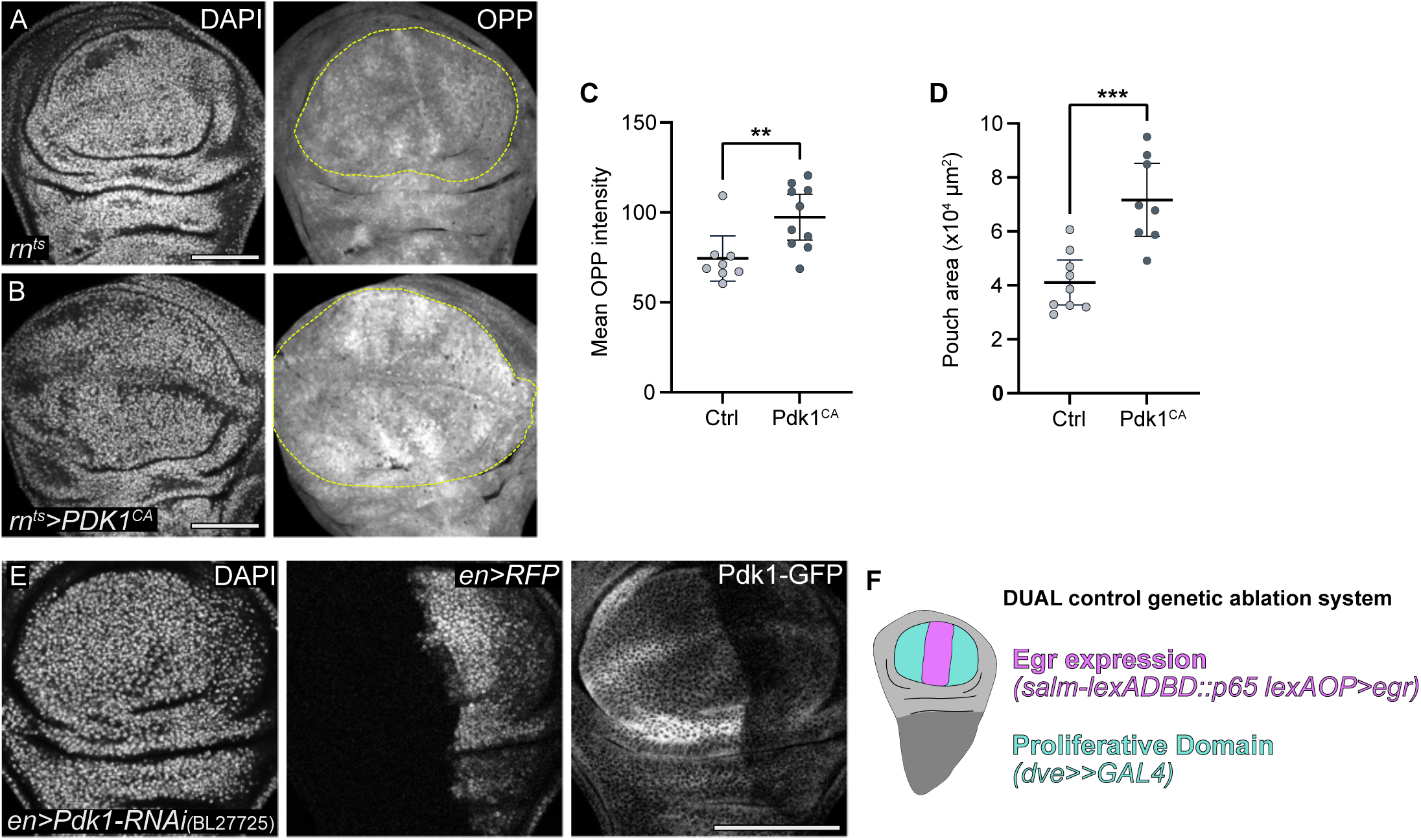
Pdk1-GFP is necessary and sufficient to drive protein translation in the regenerative domain. **A, B.** Protein synthesis visualized by OPP incorporation in control (A) and discs expressing a constitutively active UAS-Pdk1 (B) for 24 h under the control of *rn-*GAL4. Discs were stained with DAPI to visualize nuclei. The dotted white line marks the border of the pouch using tissue fold landmarks. **C.** Quantification of mean OPP intensity in the pouch of control and *Pdk1^CA^*-expressing wing discs. Mean and 95% CI are shown and statistical significance was tested using two-tailed Mann-Whitney test. p-value = 0.0062. (Control; n=8, *Pdk1^CA^*-expressing discs; n=10). **D.** Quantification of the pouch area in control and *Pdk1^CA^*-expressing wing discs. Mean and 95% CI are shown and statistical significance was tested using two-tailed Unpaired t-test. p-value = 0.0003. (Control; n=9, *Pdk1^CA^*-expressing discs; n=8). **E.** Pdk1-GFP expression in the posterior compartment of the wing imaginal disc following Pdk1 knockdown using *en-*GAL4 to test the efficiency of Pdk1-RNAi line (BL27725). *en-*GAL4 which also drives expression of UAS-RFP expression to mark the posterior compartment. DAPI stains nuclei. **F**. A simplified schematic representing DUAL control genetic ablation system (DCS). The genetically ablated region expresses *egr* under the control of *salm-lexADBD::p65 lexAOP* (marked in magenta), while *dve-*GAL4 is used to manipulate gene expression in the proliferative domain (cyan) surrounding the *egr* expressing region. Scale bar: 100 μm. Fluorescence intensities are reported as arbitrary units.

**Figures S6.**
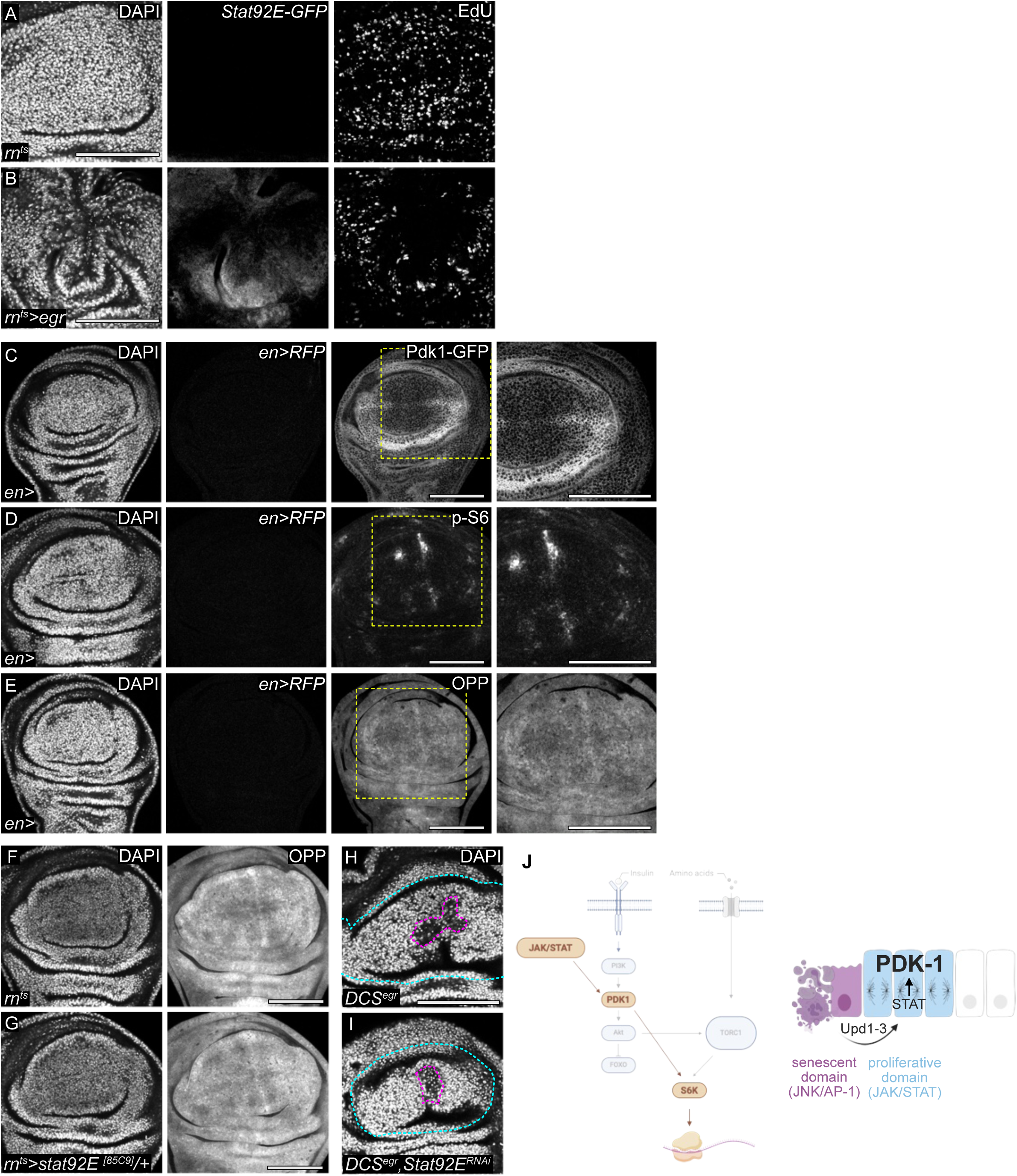
Pdk1 is regulated by JAK/STAT signaling. **A, B.** Control and *egr-*expressing discs stained for EdU to visualize DNA replication in discs also co-expressing a *Stat92E-GFP* activity reporter. Discs were also stained with DAPI to visualize nuclei. EdU intensity in *Stat92E-GFP* reporter-expressing domain was previously quantified (Jaiswal, Egert et al. 2023). **C, D, E.** Corresponding control discs to the wing imaginal discs expressing UAS-STAT92E in the posterior compartment under the control *en-*GAL4 in Fig 6**.C, E and G**. Control discs were assayed for expression of Pdk1-GFP (C), levels of p-S6 (D), and protein synthesis using OPP incorporation assays (E). *en-*GAL4 also drives expression of UAS-RFP and DAPI staining visualizes nuclei. Yellow dotted box marks the inset region. **F,G.** Protein synthesis visualized by OPP incorporation in wing discs that were either wild type (F) or heterozygous mutant for the *Stat92E^85C9^*null allele (G). DAPI staining visualizes nuclei. **H, I.** Corresponding DAPI staining of wing imaginal discs expressing the DUAL Control genetic ablation system (DCS) shown in Fig 6**.P and Q**. Early third instar (L3) larvae were heat shocked at 37°C for 75min and dissected after 40 h. A single heat shock activated both ablation (region inside the magenta dotted line) in the *salm* domain and gene manipulation in the proliferative domain via *dve-*GAL4 (*dve-GAL4* expressing region is marked with a cyan dotted line). Nuclei were visualized by DAPI staining in control (H) and *Stat92E* knockdown (I) wing imaginal disc. **J.** A model of the JAK/STAT-Pdk1-S6K axis active in proliferating cells of *egr-*expressing discs. Unpaired ligands provided by senescent-like cells in the high JNK signaling domain activating JAK/STAT in the surrounding proliferative domain, leading to Pdk1 upregulation and insulin-independent growth. Scale bar: 100 μm. Fluorescence intensities are reported as arbitrary units.

**Figures S7.**
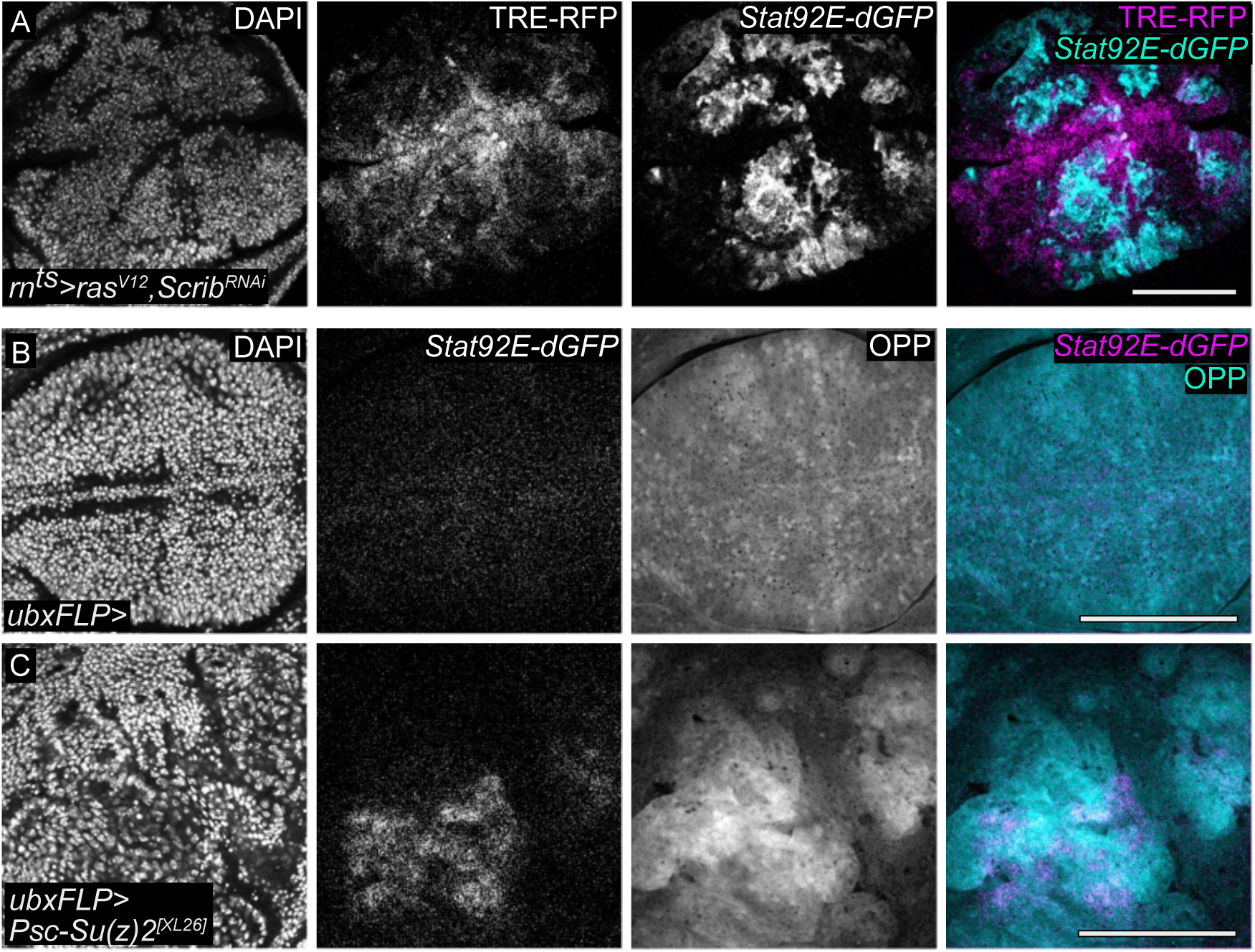
Protein translation and JAK/STAT signaling are linked in tumor growth. **A.** Wing disc expressing *Ras^V12^, scrib-RNAi* for 44h starting at Day6 AED. The disc also expresses the JAK/STAT activity reporter *Stat92E-dGFP* (cyan or grey) and JNK/AP1 reporter TRE-RFP (magenta or grey). DAPI staining visualizes nuclei. **B, C.** Control wing disc (B) and wing disc with mosaic clones mutant for the Polycomb family genes *Psc-Su(z)2^XL26^* (GFP negative cells, C). Protein synthesis visualized by OPP incorporation (cyan or grey) and JAK/STAT activity assessed using *Stat92E-dGFP* reporter (magenta or grey). DAPI staining visualizes nuclei. Scale bar: 100 μm.

**Figures S8.**
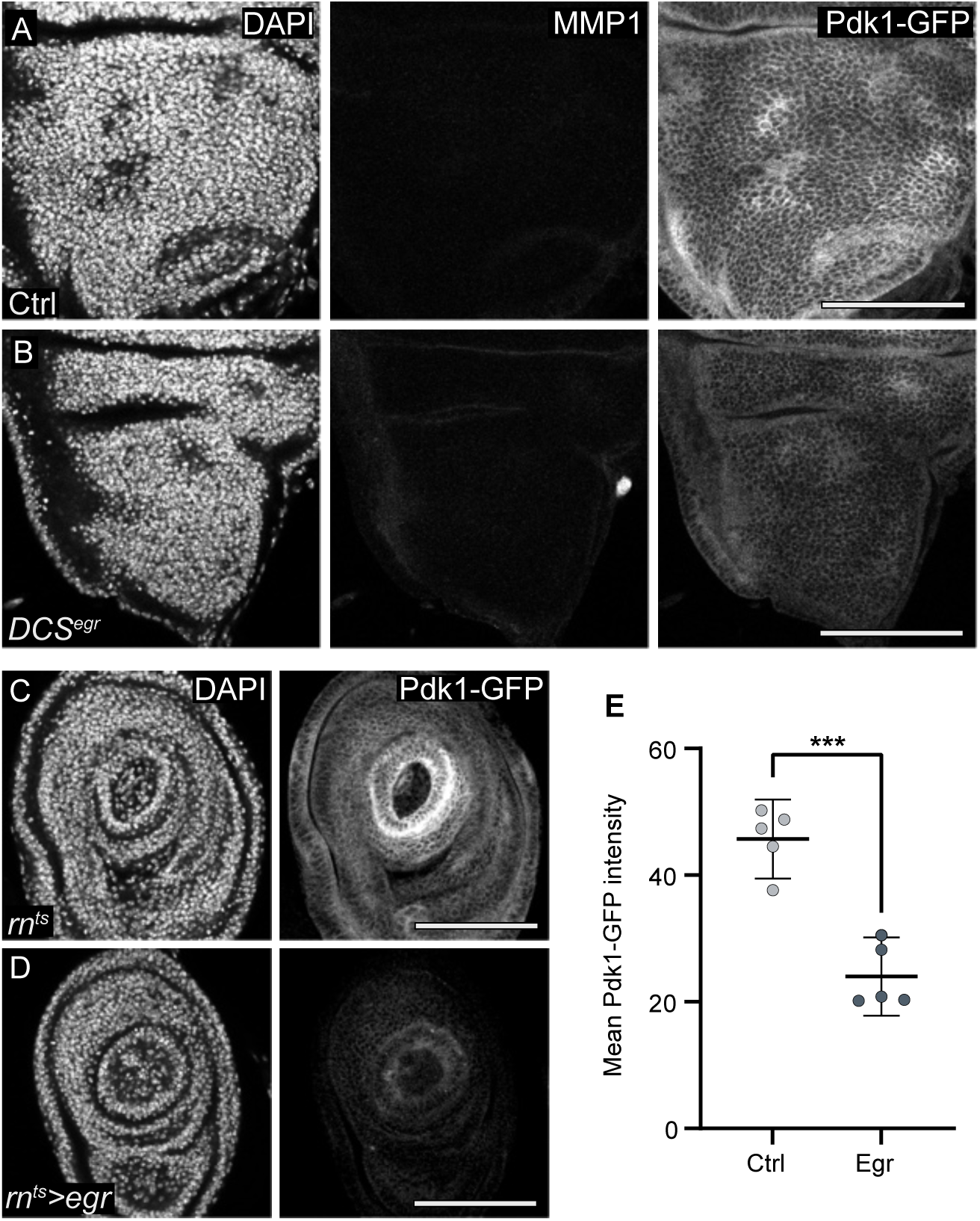
Pdk1 downregulation in peripheral discs correlates with systemic growth restriction. **A, B.** Expression of Pdk1-GFP in the notum of control (A) and *egr-*expressing discs in discs expressing the DUAL control genetic ablation system driving *egr-*expression (B). JNK/AP1 activation assessed by MMP1 staining. DAPI staining visualizes nuclei. **C, D.** Expression of Pdk1-GFP in the leg imaginal disc, dissected from larvae with control (c) or *egr-*expressing (D) wing imaginal discs expressed using the *rn*-GAL4 driver. **E.** Quantification of mean Pdk1-GFP intensity in the leg imaginal disc, dissected from larvae with control (C) or *egr-*expressing (D) wing imaginal discs. Statistical significance was tested using the two-tailed Unpaired t-test, p-value = 0.0001 (control: n=5, experiment: n=5).

**Table S1.**
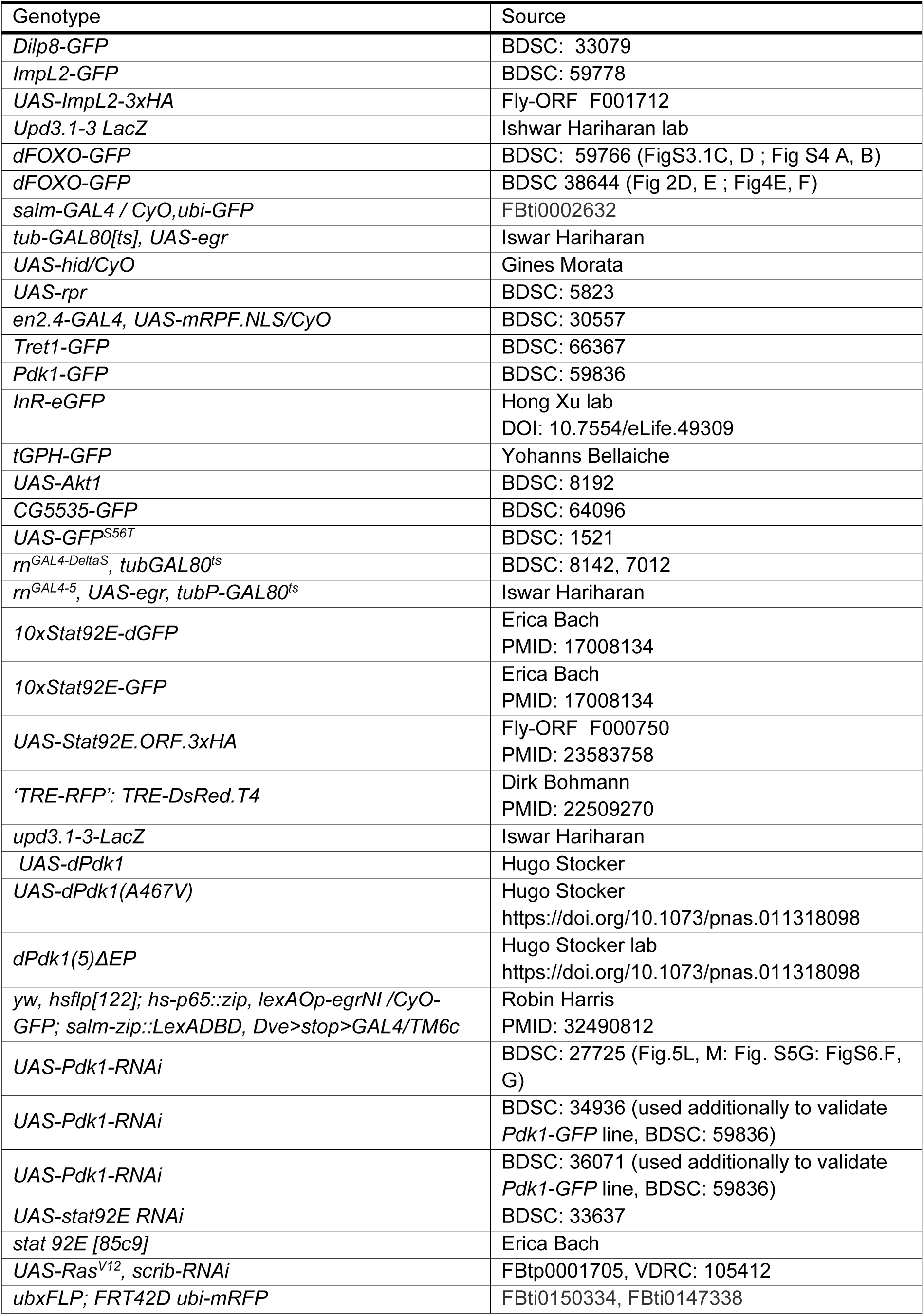
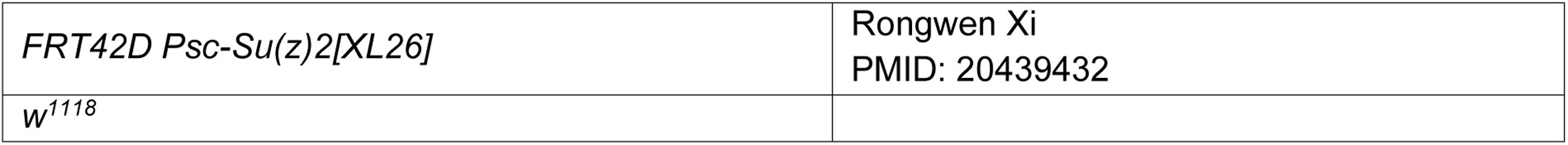
*Drosophila* strains.

**Table S2.**
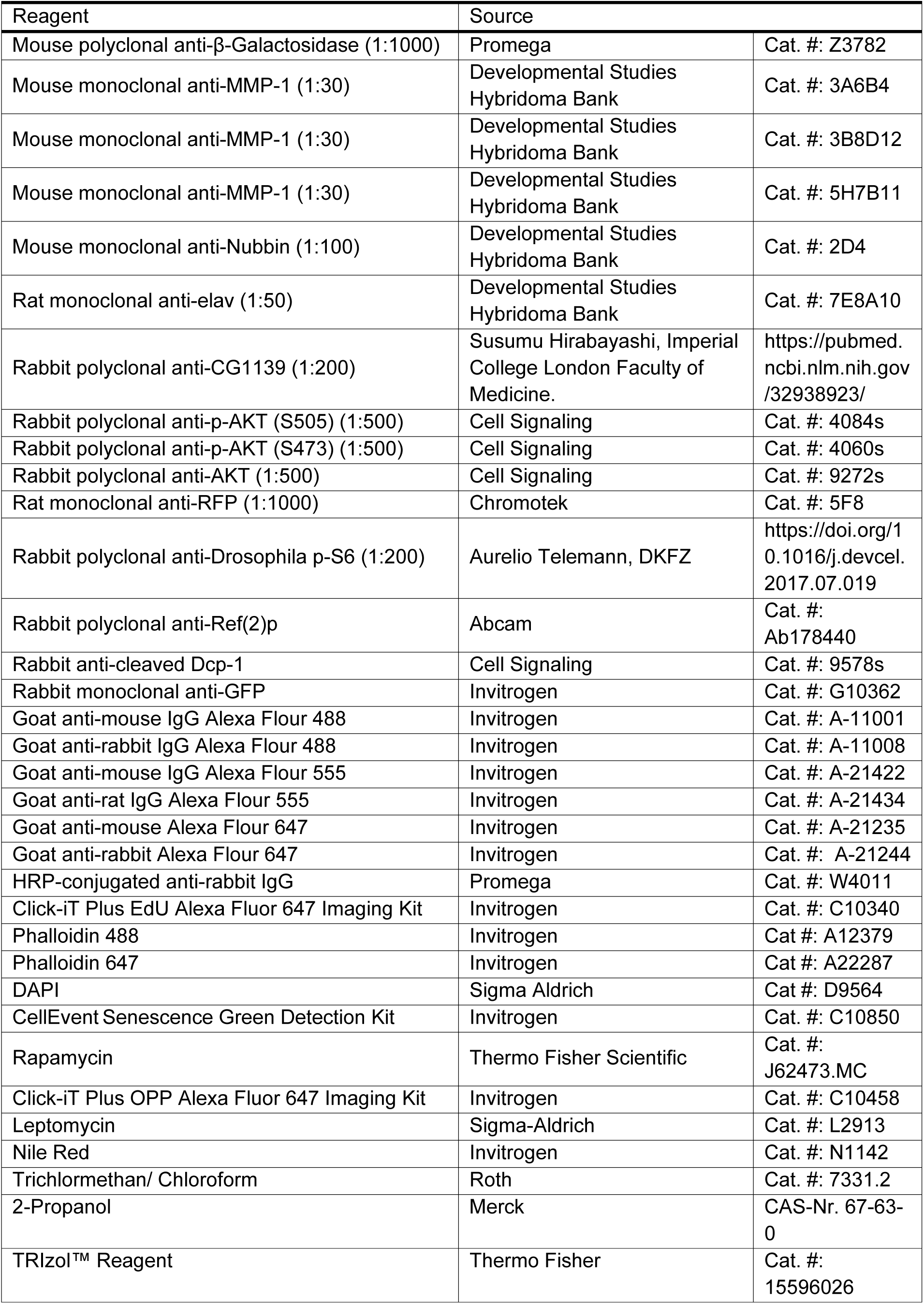
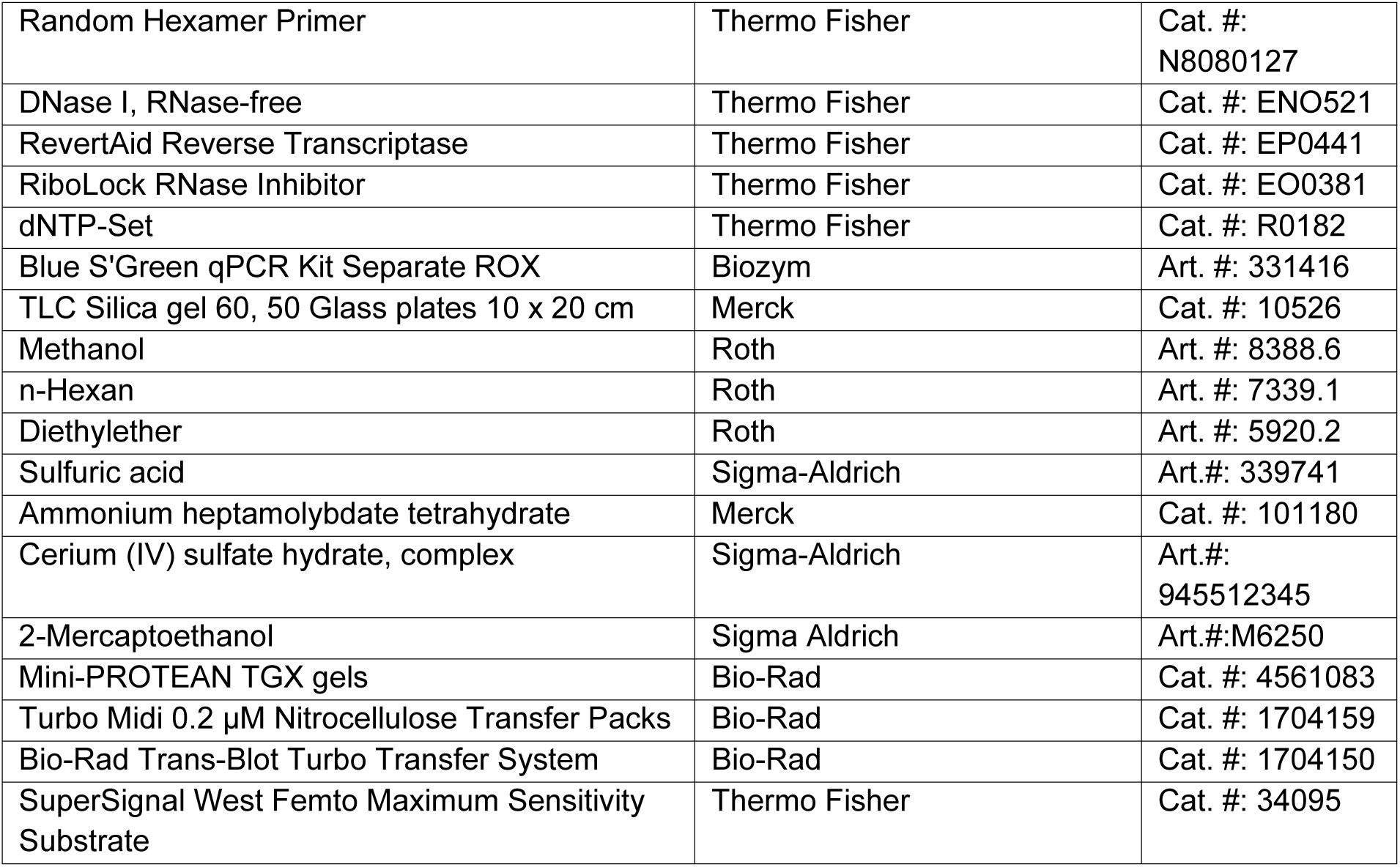
Immunohistochemistry reagents.

